# Projecting single-cell transcriptomics data onto a reference T cell atlas to interpret immune responses

**DOI:** 10.1101/2020.06.23.166546

**Authors:** Massimo Andreatta, Jesus Corria-Osorio, Sören Müller, Rafael Cubas, George Coukos, Santiago J. Carmona

## Abstract

Single-cell RNA-sequencing (scRNA-seq) has emerged as a revolutionary technology for characterizing the heterogeneity of cell populations. However, robust reference atlases that can be used to systematically interpret cellular states across studies and diseases are currently lacking. Here, we generated the first cross-study T cell atlases for cancer and viral infection and developed a novel algorithm, ProjecTILs, that enables the projection of new scRNA-seq data onto these reference atlases. ProjecTILs accurately predicted the effects of multiple perturbations, including the ablation of immunoregulatory targets controlling T cell differentiation, such as *Tox*, *Ptpn2*, *miR-155* and *Regnase-1*, and suggested novel gene programs that were altered in these cells. Moving beyond mouse models, we used ProjecTILs to conduct a meta-analysis of human tumor-infiltrating T lymphocytes (TILs), revealing a remarkable conservation of TIL subtypes between human and mouse and across cancer types. Clonotype analysis supported a model in which rare human tumor-specific effector-memory (EM)-like CD8 TILs that resemble blood-circulating EM cells, differentiate into proliferative terminal exhausted/dysfunctional effector TILs through a progenitor subtype that upregulates the exhaustion master regulator *TOX*. Our novel computational method allows exploring the effect of human and murine T cell perturbations (*e.g.* as the result of therapy or genetic engineering) in terms of reference cellular states, altered genetic programs and clonotype structure, revealing mechanisms of action behind immunotherapies and opening opportunities for their improvement.

## Introduction

In response to malignant cells and pathogens, mammals (and presumably most jawed vertebrates) mount an adaptive immune response characterized by a finely-tuned balance of several specialized T cell subtypes with distinct migratory and functional properties and metabolic lifestyles. Occasionally, however, malignant cells and pathogens escape immune control, leading to cancer and chronic infections. Antigen persistence in cancer and chronic infections profoundly alters T cell differentiation and function, leading antigen-specific cells into a collection of transcriptional and epigenetic states commonly referred to as “exhausted”^1^. The complexity and plasticity of T cells make the study of adaptive immune responses in these contexts particularly challenging.

In recent years, single-cell RNA-sequencing (scRNA-seq) enabled unbiased exploration of T cell diversity in health, disease and response to therapies at an unprecedented scale and resolution. While the presence of tumor-infiltrating T lymphocytes (TILs) in cancer lesions has been broadly associated with improved prognosis and response to immune checkpoint blockade^2,3^, scRNA-seq has uncovered a great diversity within TILs, suggesting that distinct TIL states contribute differently to tumor control and response to immunotherapies^4–6^. However, a comprehensive definition of T cell “reference” subtypes remains elusive. Poor resolution of T cell heterogeneity remains a limiting factor towards understanding the effect of induced perturbations, such as therapeutic checkpoint blockade and genetic editing, in particular when these simultaneously affect the frequencies and intrinsic features of T cell subtypes.

A major challenge towards the construction of reference single-cell atlases is the integration of gene expression datasets produced from heterogeneous samples from multiple individuals, tissues, batches and generated using different protocols and technologies. Several computational methods have been developed to correct technical biases introduced by handling experiments in batches, and to align datasets over their biological similarities^7–13^. We recently proposed STACAS^14^, a bioinformatic tool for scRNA-seq data integration specifically designed for the challenges of integrating heterogeneous datasets characterized by limited overlap of cell subtypes. This is particularly relevant for the construction of T cell atlases, where differences between datasets are not merely the result of technical variation of handling samples in different batches, but rather due to subtypes of highly variable frequency, and in many cases subtypes that are entirely missing from one or more samples as a result of study design or biological context. While whole-organism single-cell atlases are very powerful to describe the global properties of cell populations^15^, only by constructing specialized atlases for individual cell types can one achieve the level of resolution required to discriminate the spectrum of transcriptional states that can be assumed by each cell type.

A second outstanding challenge in single-cell data science is the mapping of single cells to a reference atlas^16^. The ability to embed new data points into a stable reference map has the advantage of robust, reproducible interpretation of new experiments in the context of curated and annotated cell subtypes and states. In addition, it allows the comparison of different conditions (*e.g.* pre- vs. post-treatment, or mutant vs. wild-type) over a unified transcriptomic reference landscape. In the absence of reliable reference cell atlases – and computational tools to project new data onto these atlases – researchers must rely on unsupervised, manual annotation of their data, a time-consuming and, to a certain degree, subjective process. In this work, we constructed the first cross-study atlases of tumor-infiltrating and virus-specific T cells and developed ProjecTILs, a computational framework for the projection and embedding of new scRNA-seq data into reference atlases. ProjecTILs, paired with a reference T cell atlas, enables: *i)* classification of T cells into discrete, curated subtypes, as well as *ii)* in a continuum of intermediate states along the main axes of variation of the reference map; *iii)* automated analysis of T cell perturbations; and *iv)* identification of genetic programs significantly altered in a query dataset, within individual cell subtypes, compared to the baseline reference map or to a control dataset. We demonstrated the robustness of the method by interpreting the effects of T cell perturbations in multiple model systems of cancer and infection. Finally, ProjecTILs analysis of T cell heterogeneity and clonal structure across patients, tissues and cancer types, revealed a large conservation between human and mouse TIL states and provided new insights into the differentiation of CD8 T cells in cancer.

## Results

### A cross-study reference atlas of tumor-infiltrating T cell states

With the goal to construct a comprehensive reference atlas of T cell states in murine tumors, we collected publicly available scRNA-seq data from 17 melanoma and colon adenocarcinoma tumors (see Methods). Additionally, we generated new scRNA-seq data from 4 tumor-draining lymph node samples (MC38_dLN dataset, see Methods). After data quality checks and filtering pure αβ T cells, our database comprised expression profiles of 16,803 high-quality single-T cell transcriptomes from 21 samples from six different studies.

Substantial batch effects are typically observed between single-cell experiments performed on different days, in different labs, or using different single-cell technologies. Without batch-effect correction, cells tend to cluster by study rather than by cell type (Figure 1 A). We applied the STACAS algorithm^14^ to integrate the datasets over shared cell subtypes and combine them into a unified map (Figure 1 B, Suppl. Figure S1). Unsupervised clustering and gene enrichment analysis, supported by T cell supervised classification by TILPRED^17^ (Figure 1 C), allowed annotating areas of the reference map into “functional clusters” (Figure 1 D), characterized by known gene expression signatures of specific T cell subtypes. In particular, we observed a distinct separation between CD4 and CD8 T cells, which could be further divided into subgroups. In particular, we identified a cluster of Naive-like (which might contain both naïve and central memory) CD8 T cells and a smaller cluster of Naive-like CD4 T cells, co-expressing *Tcf7* and *Ccr7* while lacking cytotoxic molecules and activation features such as *Pdcd1 and Tnfrsf9/*4-1BB; a cluster of Effector Memory-like (EM) CD8 T cells, co-expressing *Tcf7* and granzymes (most prominently *Gzmk)*, with low to intermediate expression of *Pdcd1*; an “Early activation” state of CD8 T cells, with an intermediate profile between the Naive-like and the EM-like CD8 types; a CD8 Terminally-Exhausted (Tex) effector cluster, characterized by high expression of granzymes, multiple inhibitory receptors (*Pdcd1*, *Ctla4*, *Lag3*, *Tigit*, *Havcr2*/TIM-3, etc.) and *Tox*^17,18^; a CD8 Precursor-Exhausted (Tpex) cluster, with co-expression of *Tcf7*, *Pdcd1*, *Ctla4*, *Tox* but low expression of *Havcr2* or granzymes ^17,19,20^; a cluster of CD4 Th1-like cells, expressing IFN-gamma receptor 1 (*Ifngr1*) and *Fasl*^21^; a CD4 follicular-helper (Tfh) population^21,22^, with a pronounced expression level of *Cxcr5*, *Tox* and *Slamf6*; and a cluster of regulatory T cells (Treg), identified by *Foxp3* (Figure 1 D-F). As expected, while TIL samples were enriched in Tex, Tpex and Treg subtypes, tumor-draining lymph nodes were enriched in naïve-like and follicular helper cells (Figure 1 G). Overall this new reference atlas summarizes TIL diversity using nine broad cell subtypes with distinct phenotypes, functions, metabolic lifestyles and preferential tissue distributions, and strongly supported by experimental evidence in murine models. An interactive interface of the TIL atlas, allowing the exploration of T cell subtypes and gene expression over the reference map can be accessed at http://TILatlas.unil.ch

**Figure 1.**
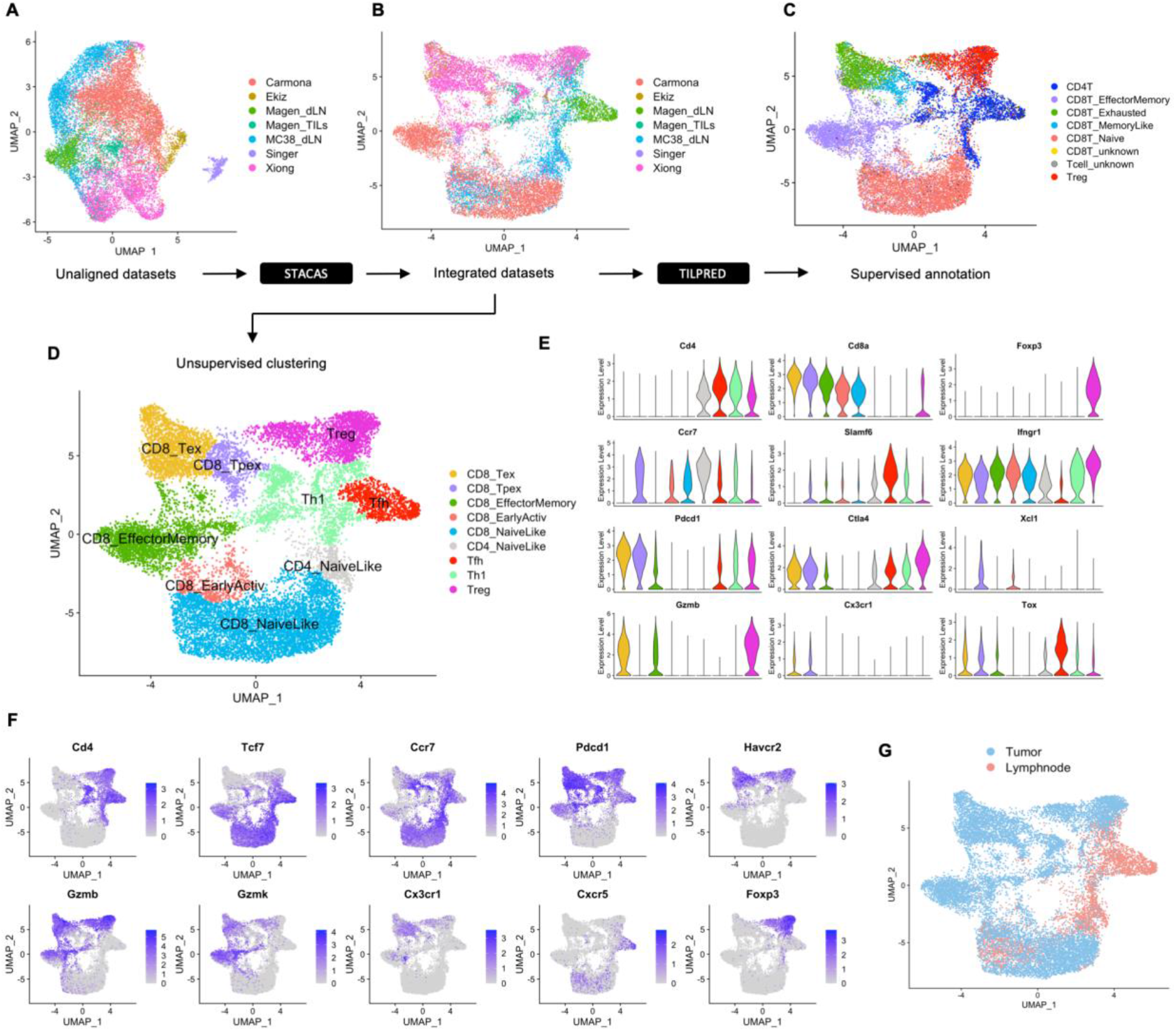
Building a reference map of TIL transcriptomic states. **A)** Uniform Manifold Approximation and Projection (UMAP) plots of single-cell transcriptomic profiles from different studies, before batch-effect correction (*i.e.* unaligned datasets); **B)** Same plot for integrated datasets after STACAS alignment: successful dataset integration mitigates batch effects while preserving biological differences between T cell subtypes; **C)** Supervised T cell subtype classification by TILPRED shows that, after alignment, cells cluster mainly by cell subtype rather than by dataset of origin; **D)** Unsupervised clusters were annotated as nine functional states based on TILPRED prediction, as well as by **E)** average expression of marker genes in each cluster and by (**F**) single-cell expression of key marker genes over the UMAP representation of the map; **G)** Reference atlas colored by tissue of origin (tumor and draining lymphnode). An interactive reference TIL atlas can be explored online at http://TILatlas.unil.ch

### ProjecTILs enables accurate projection of scRNA-seq data onto reference atlases

In order to enable interpretation of new datasets in the context of reference T cell subtypes, we developed ProjecTILs, a novel method for the projection of scRNA-seq data onto a reference atlas. The essential input to ProjecTILs is the single-cell expression matrix of the query dataset, *e.g.* in UMI counts or TPMs, where rows represent the genes and columns represent the individual cells. The pre-processing steps (Figure 2 A) normalize scRNA-seq data using a log-transformation (if provided non-normalized) and filter out non-T cells (see Methods). In order to reduce batch effects between the query and the reference map, the STACAS/Seurat integration procedure^14^ is used to align the query to the reference, and in this way correct the expression matrix of the query dataset (Figure 2 B, see Methods). The corrected query matrix can then be projected onto reduced-dimensionality representations (e.g. PCA, UMAP) of the reference, effectively bringing them into the same reference space. To this end, the algorithm computes the PCA rotation matrix of the reference map, which contains the coefficients to linearly transform gene expression into PCA loadings (i.e. the eigenvectors and their relative eigenvalues); the same PCA rotation matrix is then also applied to the query set (Figure 2 C). Likewise, the UMAP transformation (allowing the computation of UMAP coordinates from PCA loadings) is applied to the query set to project it into the original UMAP embedding of the reference map. Note that while the expression counts of the query set and their embedding into reduced spaces are modified by the alignment procedure, those of the reference map are not, and its low-dimensional visualizations remain unaltered; different query sets, or experiments containing different conditions can therefore be compared over the same reference map. After projection, a nearest-neighbor classifier predicts the subtype of each query cell by a majority vote of its annotated nearest neighbors (either in PCA or UMAP space) in the reference map. Benchmarking ProjecTILs by cross-validation experiments showed a high accuracy (>90%) both for the classification (Suppl. Figure S2) and projection tasks (Suppl. Figure S3, and Methods).

**Figure 2:**
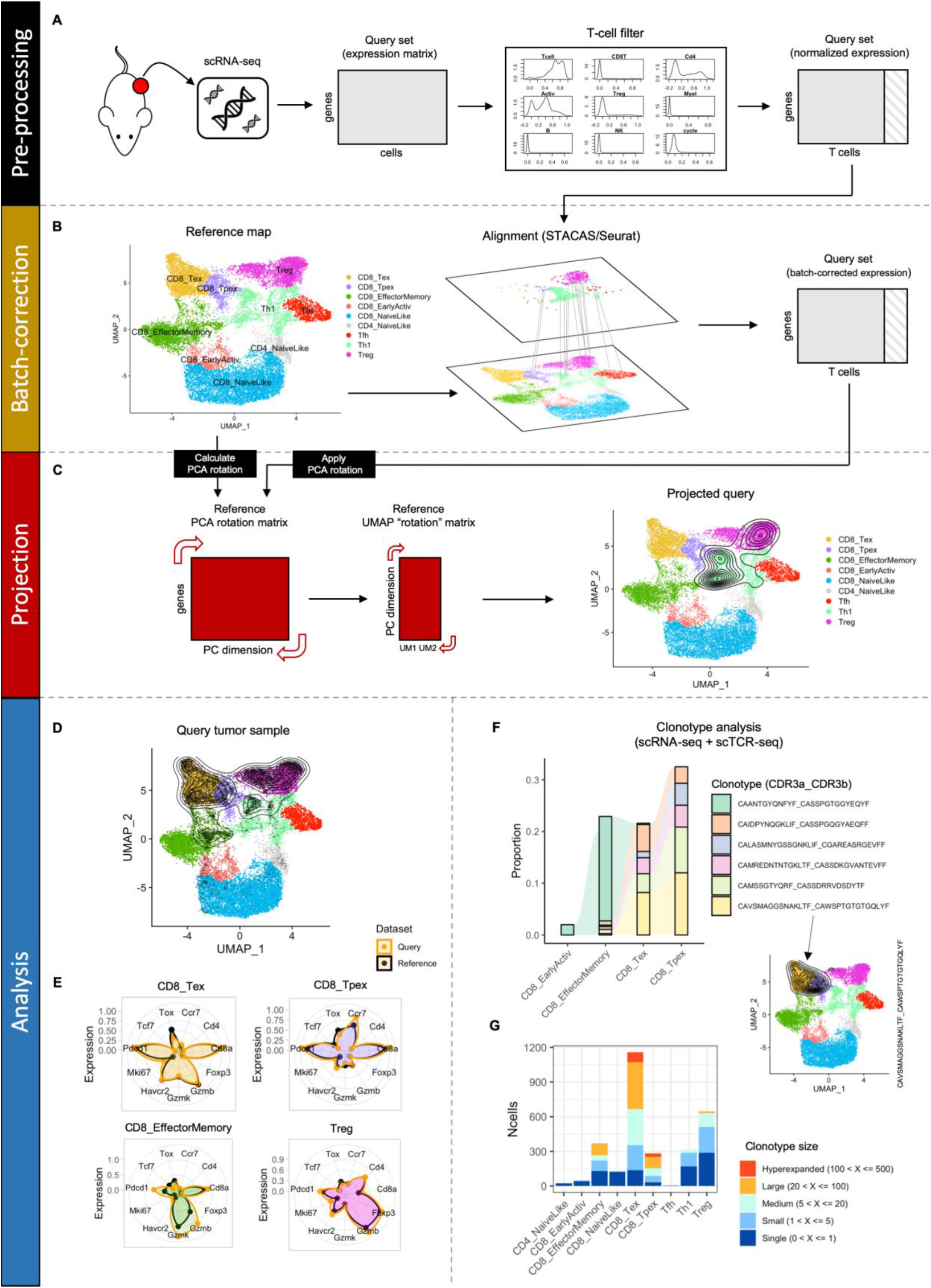
The ProjecTILs analysis workflow. **A)** The essential input to ProjecTILs is a query dataset in the form of a gene expression matrix. Pre-processing steps include data normalization and filtering of non-T cells. **B)** The normalized, filtered gene expression matrix is aligned to the reference map using STACAS, to bring the query data into the same scale as the reference map. **C)** The PCA rotation matrix and UMAP transformation calculated on the reference map are applied to the query set, effectively embedding it into the same space of the reference map, and allowing their direct comparison and joint visualization. **D-G)** Projection example of tumor-infiltrating T cells from Xiong *et al.*^23^: **D)** Predicted coordinates of the projected query in UMAP space as density contours. **E)** Gene expression signature of query cells (orange) and reference cells (black) for four selected T cell subtypes. **F)** Proportion of cells across CD8 T cell subtypes for the six largest clonotypes, highlighting overlap between related T cells subtypes; the inset shows the projection of the single largest clonotype (CDR3a:CAVSMAGGSNAKLTF CDR3b:CAWSPTGTGTGQLYF). **G)** Clonotype size (single to hyperexpanded) of T cells across the ProjecTILs predicted subtypes.

As an illustrative application of ProjecTILs to analyze a query scRNA-seq dataset, we projected onto the reference TIL atlas a dataset of tumor-infiltrating T cells (TILs) from the study by Xiong *et al.*^23^. The density distribution of projected cells reveals that the majority of TILs are predicted to be CD8 exhausted, precursor exhausted and Tregs, and to a lesser extent CD8 EM cells (Figure 2 D). Within these compartments, the expression profiles of query cells and reference cells display very high correspondence, confirming the accuracy of the projection (Figure 2 E). When single-cell T cell receptor sequencing (scTCR-seq) data are available, ProjecTILs interacts with scRepertoire^24^ to allow exploring the clonal diversity and distribution across the T cell subtypes of the reference atlas (see also https://carmonalab.github.io/ProjecTILs_CaseStudies/Xiong19_TCR.html). Of the six largest CD8 T clonotypes identified among these TILs, five appear to be shared across the Tex and Tpex subtypes, while one is almost exclusive to the EM compartment (Figure 2 F), in agreement with previously reported TIL clonal structure in mice^17^. Globally, Tex and Tpex show the largest level of clonal expansion, followed by EM and Tregs, while the remaining subtypes are mostly composed of single or poorly expanded clones (Figure 2 G).

To explore the states of tumor-reactive TILs, we projected onto the reference murine TIL map a dataset of tumor-specific CD8 T cells isolated by tetramer staining from untreated B16 melanoma tumors expressing chicken ovalbumin (OVA), from the study by Miller *et al.*^25^. Consistently, ProjecTILs assigned the great majority (90.2%) of tumor-specific cells to the Tex subtype, while small fractions were assigned to the Tpex (4.4%) and EM-like (3.9%) compartments (Suppl. Figure S4 A-B). The expression profile of Tex cells matches well with the reference map for a panel of marker genes, with a pronounced overexpression of *Gzmb* and the proliferation marker *Mki67* (Suppl. Figure S4 C), as expected by antigen-induced activation^26,27^. ProjecTILs allows visualizing an additional dimension on the z-axis together with the 2D UMAP representation. In this case, we chose to plot the cell cycling score calculated by TILPRED, which reveals a striking proliferative signal for the cells in the query dataset (Suppl. Figure S4 D). Taken together, these results are in agreement with experimental observations and well compatible with the notion that tumor-specific CD8 T cells, as a result of continued antigenic stimulation in the tumor, become enriched in a highly expanded, terminally exhausted state^25,28^.

### ProjecTILs reveals altered states and gene programs following T cell perturbations

CD8 T cells depend on *miR-155* to acquire anti-tumoral or anti-viral effector functions^29,30^. To interpret the impact of *miR-155* deficiency on the TIL landscape, we submitted to ProjecTILs the data from Ekiz *et al.*^31^, which consist of total immune (CD45+) single cells from untreated B16 melanoma tumors growing in microRNA 155 (*miR-155*) T cell conditional knock-out (KO) mice, as well as in wild type (WT) mice. Projecting these data onto the reference TIL atlas revealed that in WT mice, the majority of TILs correspond to CD8 Tex, with smaller populations of other cell types. Conversely, CD8 TILs from *miR-155* KO mice were mostly projected to the naive-like compartment (Figure 3 A). Moreover, while T cells accounted for 16% of the total immune infiltrate in WT mice, they were reduced to 8% in KO mice. Consistently with the predicted change in the dominant T cell phenotype from exhausted to naive-like upon *miR-155* KO, the expression of activation markers (e.g. *Tnfrsf9* and *Ifng*) was reduced in the KO mice, while memory/naive markers such as *Tcf7* and *Ccr7* were overexpressed (Figure 3 B). The KO TILs also scored lower in terms of the cycling signature (Figure 3 C). Altogether, these results are consistent with the critical role of *miR-155* for T cell activation and differentiation. Unable to differentiate and acquire effector functions, *miR-155* KO CD8 TILs fail to control tumor growth^31^. In brief, by comparing the changing landscape and cell subtypes distribution of the KO compared to the WT TILs, ProjecTILs provides a straightforward interpretation of the effect of this genetic alteration, all in the context of annotated, reference T cell subtypes.

**Figure 3:**
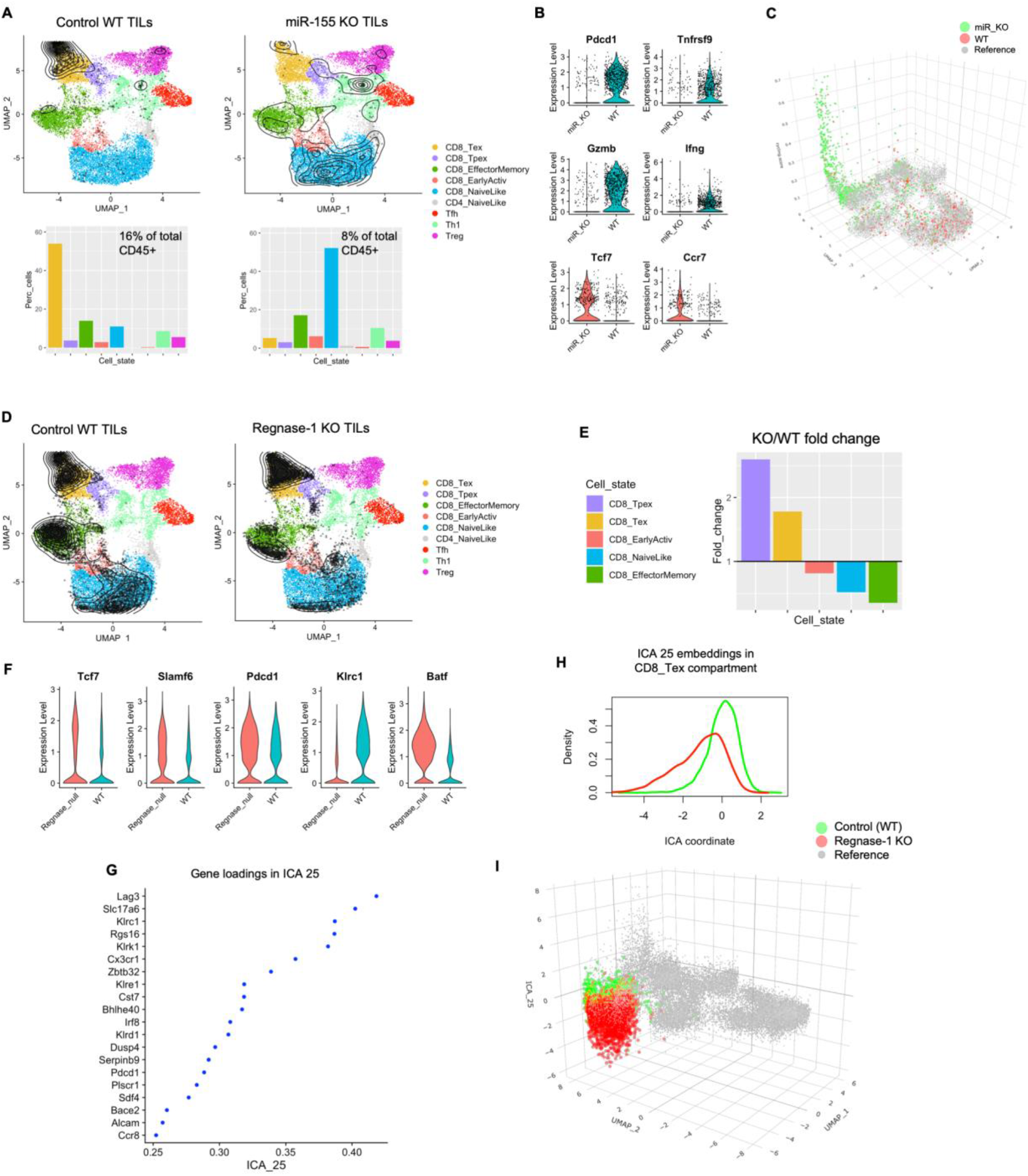
ProjecTILs reveals the effect of genetic perturbations on T cell transcriptomes and phenotypes. **A-C)** ProjecTILs analysis of the tumor CD45+ scRNA-seq data by Ekiz *et al.*^31^: **A)** WT and *miR-155* KO TILs projected on the reference atlas (black points and density contours) and barplots depicting percentage of cells projected in each T cell subtype for the two conditions. T cells constituted 16% and 8% of the CD45+ cells for the WT and *mi*R-155 KO samples, respectively; **B)** Violin plots showing expression of activation and cytotoxicity (*Pdcd1*, *Tnfsf9*/4-1BB, *Gzmb*, *Ifng*) and naïve/memory (*Tcf7*, *Ccr7*) markers; **C)** Cell cycling score represented on the z-axis of the UMAP for the reference map of WT cells and *miR-155* KO cells. **D-I)** ProjecTILs analysis of the scRNA-seq data by Wei *et al.*^32^: **D)** Cells projection on reference TIL atlas (similar to A); **E)** Fold-change in the *Regnase-1* KO compared to WT for each TIL subtype containing at least 50 cells; **F)** Global expression level for selected genes in the *Regnase-1* KO versus the WT; **G)** Top 20 driver genes in terms of gene loadings for the transcriptional program ICA 25, the most discriminant dimension between the WT and the *Regnase-1* KO; **H)** Distribution of ICA 25 coordinates for cells in the WT (green) versus *Regnase-1* KO (red) samples, as can be also visualized (**I**) by plotting these values on the z-axis of the UMAP plot.

While low-dimensional representations such as the UMAP are useful to summarize the most prominent transcriptional features that discriminate distinct T cell subtypes, they are often not sufficient to fully capture the heterogeneity of cell transcriptomes. With the goal to provide more resolution to the analysis of T cell states, and in particular to identify gene programs that are shared by multiple cell subtypes, we decomposed the reference atlas into 50 dimensions using Independent Component Analysis (ICA) (see Methods). Because ICA finds a representation of the data where the dimensions share minimal mutual information, it can be useful to separate different gene modules and transcriptional programs within complex gene expression datasets^33^ and provides a complementary description to the UMAP representation – which in turn is built over a PCA reduction of the transcriptomic space. In order to interpret the biological relevance of the ICA components in the reference TIL atlas, we investigated their correlation with annotated molecular signatures from the mSigDB database^34^ and observed several modules associated with key cellular pathways (Suppl. Figure S5). For example, the component ICA 33 was driven by several genes associated with hypoxia (e.g. *Tpi1*, *Pkg1*, *Ldha*, *Slc2a1*); ICA 40 was rich in E2F targets (*Mcm2* to *Mcm7*, *Dut*, *Cdc6*), indicating a module of key regulators of cell cycle progression; ICA 26 contained multiple genes involved in cytotoxicity, such as *Prf1* (encoding perforin 1) and several granzymes (*Gzme*, *Gzmc*, *Gzmb*); and ICA 37 was strongly associated with response to interferons (e.g. *Ifit1*, *Ifit3*, *Rsad3*) (Suppl. Figure S6). Critically, some ICA components appeared to affect mainly specific T cell subtypes or regions in the context of our reference map (*e.g* ICA 43 and 45), while others captured features of multiple subtypes/regions (*e.g.* ICA 33 and 37, Suppl. Figure S7).

Importantly, we can compare a projected query dataset with the reference map – or two query conditions between themselves – in their ICA representations, and identify components in which the query dataset deviates from the reference. ICA dimensions where the two sets differ significantly suggest gene modules that are up- or down-regulated in the query set, and may aid the biological interpretation of experimental observations. As an illustrative example of this approach, we re-analyzed the scRNA-seq from Wei *et al.*^32^. In this study, authors show that ablation of *Zc3h12a* (which encodes Regnase-1) in CD8 T-cells improved the therapeutic efficacy of adoptively transferred, tumor-specific cells in mouse models of melanoma and leukemia. Projection of the Regnase-1-null CD8 TILs and WT counterparts into our TIL reference atlas showed that cells from the two conditions occupied similar CD8 regions of the map (Figure 3 D). However, Regnase-1-null CD8 T cells displayed a 3-fold relative enrichment in Tpex and a 1.8-fold enrichment in Tex compared to WT T cells (Figure 3 E). Consistently, Regnase-1-null TILs showed an overall increased expression of *Tcf7*, *Slamf6* and *Pdcd1* (Figure 3 F). Ablation of Regnase-1 favored acquisition of the Tpex state, explaining the increased T cell persistence reported by Wei et al. However, T cell-mediated tumor control additionally requires enhanced effector functions. To investigate whether Tex Regnase-1-null cells displayed altered gene programs compatible with an enhanced effector state, in addition to their increased relative abundance, we applied ProjecTILs discriminant ICA components analysis. Interestingly, ICA discriminant analysis of Tex cells between the two conditions revealed that the most significant deviation was in ICA 25 (KS test statistic = 0.612, p-val=0), a component driven by the checkpoint molecules *Lag3* and *Klrc1* (Figure 3 G-H). Visualizing the ICA 25 coordinates of the Regnase-1-null cells on the z-axis of the UMAP plot, we clearly observe that this component is highly down-regulated compared both to the WT data and the reference map (Figure 3 I). Accordingly, *Klrc1* expression was lower in Regnase-1-null cells (Figure 3 F). While analyzing the expression levels of key marker genes can suggest global patterns of alteration due to genetic perturbations (Figure 3 F), only by comparing gene programs in each cell subtype individually across conditions can one pinpoint the molecular programs affected by the perturbation while avoiding confounding effects due to global shifts in relative proportions of T cell subtypes. Importantly, a similar distribution of predicted cell subtypes and significant down-regulation of ICA 25 could be detected even in the absence of a control sample, by direct comparison of the Regnase-1-null cells to the reference map (Suppl. Figure S8).

### ProjecTILs beyond cancer: interpreting T cell perturbations in the context of acute and chronic infections

While ProjecTILs was originally conceived to study the diversity of TILs, it can be readily applied to any other biological context for which a reference atlas can be constructed. A prominent example is the lymphocytic choriomeningitis virus (LCMV) infection model, one of the best-studied models of acute and chronic viral infection. In order to construct a reference atlas for viral infection models, we collected scRNA-seq data of virus-specific CD8 T (P14) cells^35–37^ from three different studies, consisting of single-cell gene expression measurements at different time points for acute and chronic LCMV infection (Figure 4 A). Alignment by STACAS (Figure 4 B) was followed by unsupervised clustering (Figure 4 C, see Methods). By inspecting the gradients of gene expression across the UMAP representation of the atlas (Figure 4 D), as well as the average expression of a panel of marker genes in the different unsupervised clusters (Figure 4 E), we annotated seven functional clusters: Effector Early, Effector Intermediate, Effector Cycling, Memory Precursor, Short-Lived Effector Cells (SLEC), Precursor Exhausted (Tpex), and Exhausted (Tex) (Figure 4 C). Cells from early acute infection (day 4.5) were mostly located in the Early, Intermediate and Cycling Effector areas, as well as in the Memory Precursor subtype; however, at a later time point (day 7.5) their distribution of cell subtypes shifted towards the SLEC subtype (Figure 4 F). Similarly, early chronic infection (day 4.5) was characterized by Effector T cell types, nearly indistinguishable from the acute cells at the same time point; but as the infection progressed (days 7.5 and 30) their subtype distribution diverged from the acute infection towards Tpex and Tex subtypes (Figure 4 G) with a non-persistent wave of SLEC-like cells at an intermediate time point (day 7.5).

**Figure 4:**
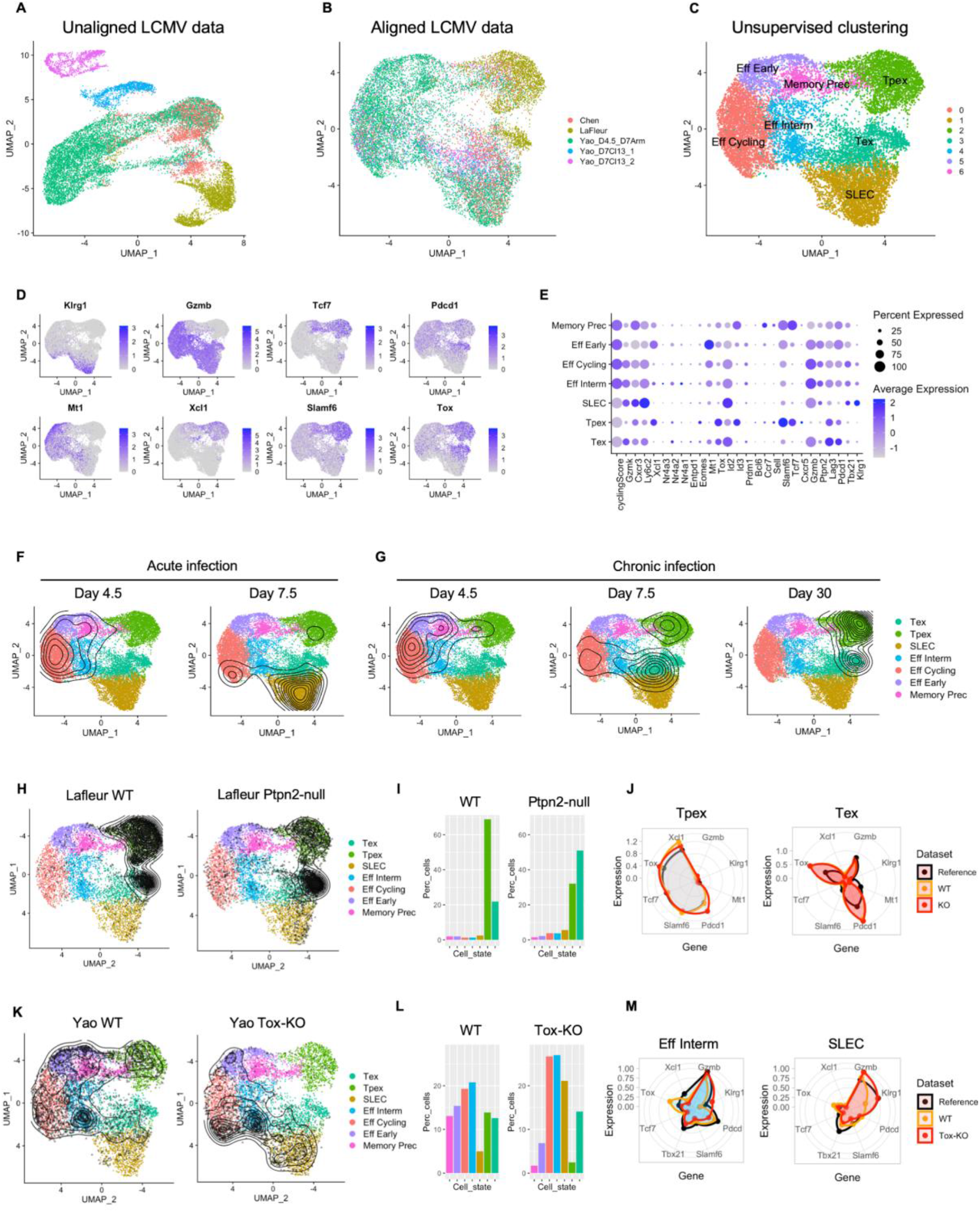
Beyond TILs: a reference atlas of virus-specific CD8 T cells during acute and chronic infection. **A)** Unaligned datasets of lymphocytic choriomeningitis virus (LCMV)-specific CD8 T (P14) cells during infection show pronounced batch effect, which **B)** can be mitigated by STACAS alignment. **C)** Unsupervised clusters were annotated to seven functional clusters by examining **D)** the gradient of expression and **E)** the average expression of marker genes by cluster (*i.e.* Memory Precursors; “Early”, “Cycling”, “Intermediate”, and short-lived (“SLEC”) effectors; Precursor Exhausted (Tpex) and Terminal exhausted (Te) CD8 T cells). **F)** Density of cells across the map at two time points in acute infection and **G)** at three different time points in chronic infection. **H-J)** Analysis of *Ptpn2* KO versus control (WT) using the data by Lafleur *et al.*^36^: **H)** ProjecTILs projection of WT and *Ptpn2* KO cells onto the infection reference map; **I)** Predicted percentage of cells for each T cell subtype; **J)** Normalized average expression for selected markers in the reference map, in WT and *Ptpn2* KO cells. **K-M)** Analysis of *Tox* KO versus control (WT) using the data by Yao *et al.*^37^: **K)** Projection in UMAP space by ProjecTILs for the WT and *Tox* KO samples; **L)** Predicted percentage of cells for each T cell type; **M)** Normalized average expression for selected markers in the reference map, in WT and *Tox* KO cells.

With this reference virus-specific CD8 T cell atlas in hand, we proceeded to project new datasets to study the effect of genetic alterations on CD8 T cells during viral infection. The phosphatase PTPN2 has been recently proposed as an attractive immunotherapeutic target to enhance T cell cytotoxicity in chronic infection and cancer^36,38^. Automated ProjecTILs analysis of co-transferred *Ptpn2*-KO and control P14 cells at day 30 after LCMV infection and CD4 depletion^36^ revealed that the large majority of cells, both *Ptpn2*-KO and WT, were projected either in the Tpex or Tex clusters (Figure 4 H). Importantly, while the Tex compartment constituted only about 20% of the WT T cells, it amounted to over 50% of *Ptpn2*-KO cells (Figure 4 I). Moreover, analysis of the average expression of key marker genes confirmed a good agreement between Tpex expression profiles (for control, KO and reference subtype), as well as overexpression of *Pdcd1* and *Tox* in the *Ptpn2*-KO cells (Figure 4 J). Therefore, ProjecTILs automated analysis supported the original observations that in chronic infection, *Ptpn2* deletion promotes differentiation of Tpex into Tex, which translates into a higher number of effector cells, improving – at least transiently – viral and tumor control^36^.

As a second example of projection, we applied ProjecTILs to study the effect of deleting the transcription factor *Tox* in virus-specific CD8 T cells during chronic viral infection. Projection of the data from Yao *et al.*^37^ revealed that *Tox*-KO cells had a dramatic alteration in subtype composition compared to WT controls (Figure 4 K). In particular, it showed a large increase in the fraction of SLECs, at the expense of Memory Precursors and Precursor Exhausted cells (Figure 4 L). Analysis of marker profiles confirmed that the *Tox*-KO cells classified as SLEC express high levels of *Klrg1* (Figure 4 M). These results are consistent with previous studies that demonstrated that TOX is required for the establishment of the exhaustion CD8 T cell program^18,37,39–41^.

Finally, we projected P14 cells from CD4-depleted (or isotype control) chronically infected mice from Kanev *et al.*^42^. We observed that anti-CD4 antibody-treated mice had a dramatic shift in their T cell subtype composition from short-lived effectors (SLEC) to Precursor Exhausted (Tpex) compared to control mice, while the proportion of Tex remained similar (Suppl. Figure S9 A-B). This effect is in agreement with the conclusions of the original study, which found that in chronic infections, progenitor cells (Tpex) are unable to properly differentiate into effector cells in the absence of CD4 T cell help^42^, and therefore “accumulate” in this state. We could also confirm that CD4 depletion had an effect on increasing *Pdcd1* expression, especially in Tex cells (Suppl. Figure S9 C). These data were generated using the SCRB-seq protocol^43^, a type of sequencing that was not included in the reference LCMV atlas. Therefore, this example also highlights the robustness of ProjecTILs to accurately project, and enable the correct interpretation of, single-cell data across multiple sequencing platforms.

### ProjecTILs beyond mouse: projection of scRNA-seq data from cancer patients onto a mouse atlas reveals a strong conservation of T cell subtypes across species

Mouse models are essential to gain mechanistic insights into tumor immune responses. Yet, precise definition of TIL subtypes conservation between human and mouse has remained elusive. Here we asked whether the T cell states described in human tumors have a clear mapping to mouse TIL subtypes. Using orthologous genes between the two species, we applied ProjecTILs to analyze human TIL scRNA-seq data from 30 cancer patients from two cohorts (melanoma cohort of Li *et al.*^5^ and basal cell carcinoma cohort of Yost *et al.*^44^) in the context of the reference murine TIL atlas (see Methods). Projected TILs from individual patients broadly distributed over the reference murine atlas (Figure 5 A, Suppl. Figure S10).

**Figure 5:**
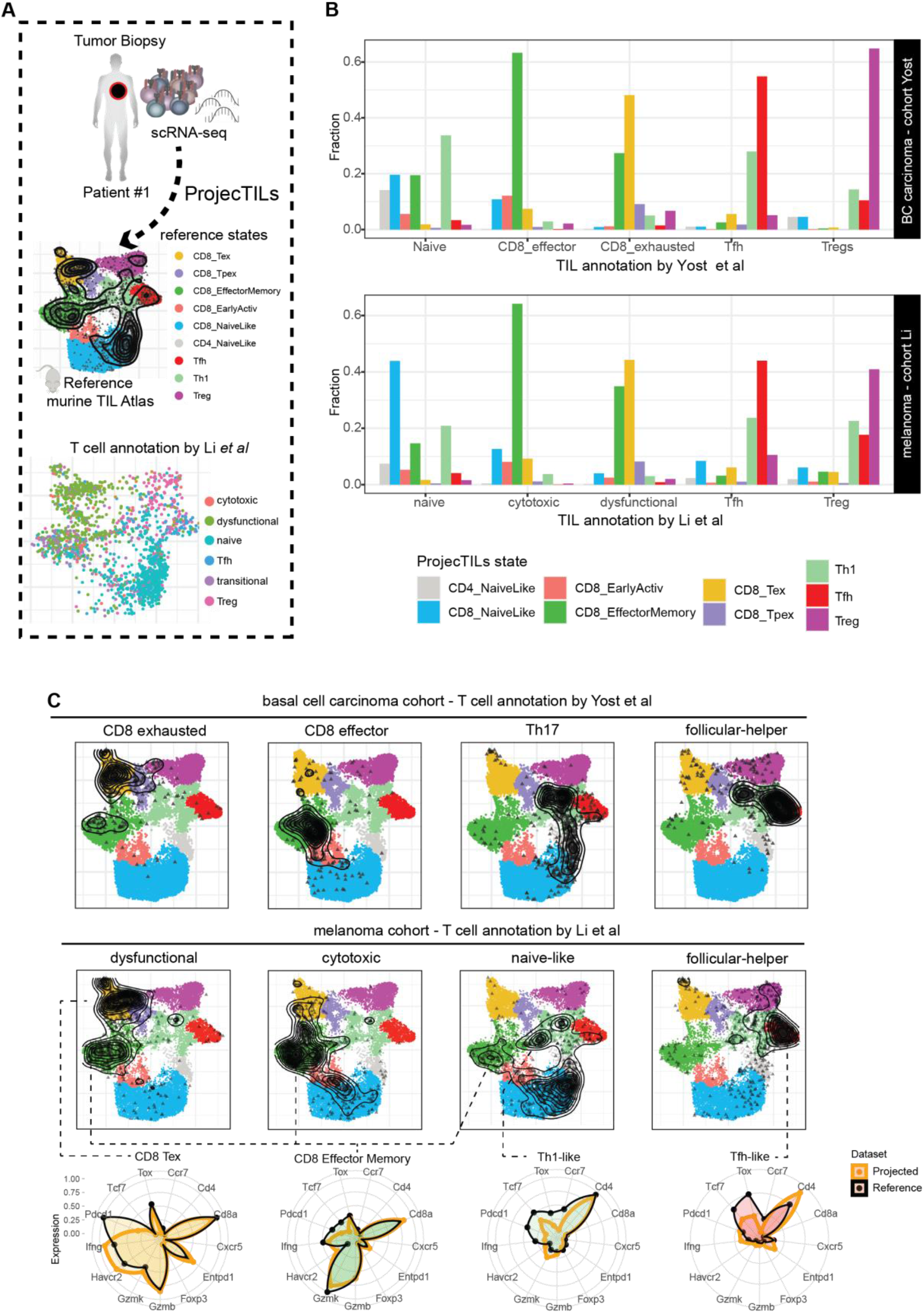
Accurate classification of human TIL states by projecting cancer patients’ transcriptomes on a reference murine atlas. **A)** scRNA-seq data from patients’ biopsies were analyzed using ProjecTILs in human-mouse orthology mode. Below, UMAP projection for TILs from one subject, colored by annotation according to Li *et al.* Projections for other subjects are available in Suppl. Figure S10. **B)** Fraction of cells classified in different subtypes by ProjecTILs compared to main original annotations by Yost *et al.* or Li *et al.* (complete annotation in Suppl. Figure S11). **C)** UMAP projections of cell subsets defined according to TIL state annotations by Yost *et al.* (e.g. exhausted, effector) or Li *et al.* (e.g. dysfunctional, cytotoxic). Radar plots display representative expression profiles of cells classified in the reference states for T cell marker genes.

ProjecTILs T cell subtype classification showed a remarkable correspondence with the T cell annotations assigned by the authors of the original studies (Li *et al.*^5^ and Yost *et al.*^44^). For example, human TILs originally defined as Treg, follicular-helper, naïve or T-helper were largely projected to the corresponding subtypes on the murine reference atlas; the “CD8 effector” (Li *et al.*) and “CD8 cytotoxic” (Yost *et al.*) cells were projected to the reference effector memory (EM) subtype; the “CD8 exhausted” cells (Yost *et al.*) and “CD8 dysfunctional” cells (Li *et al.*), were mostly projected to the reference exhausted subtype (Figure 5 B, Suppl. Figure S11). Surprisingly, a significant fraction of the cells annotated by the authors as “exhausted/dysfunctional” were projected on the EM state of the murine atlas. Further examination revealed that these cells displayed a clear effector-memory gene profile (i.e. high *GZMK, GZMA* and *GZMB* expression) but lacked markers of exhaustion, such as *TOX*, *ENTPD1*, *HAVCR*, *PDCD1* (Figure 5 C), indicating that they were correctly projected on the EM reference subtype. Similarly, “exhausted/dysfunctional” cells classified by ProjecTILs as Tex did closely match the expression profile of the Tex reference state. Overall, the correspondence between ProjecTILs classification and the human TIL state annotations defined in two different studies and cancer types is remarkable. This points to a large conservation between human and mouse TIL states.

To further validate the accuracy of ProjecTILs, we analyzed biopsies taken at baseline from a third cohort (Sade-Feldman *et al.*^6^), consisting of 19 melanoma patients that were classified as responders (R) and non-responders (NR) to checkpoint blockade. In agreement with the authors observations, ProjecTILs revealed that *TCF7*-high TIL subtypes were enriched in responders vs non-responders (Suppl. Figure S12 A). Intriguingly, these subtypes corresponded to naïve-like CD8 and CD4 cells, and not to TILs displaying markers of tumor-reactivity, such as PD-1^45^. A similar subtype bias was found between metastatic lymph nodes and tumors, irrespectively of the patients’ responsiveness to immunotherapy (Suppl. Figure S12 B). This prompted us to analyze in more depth the defining features of tumor-reactive human TILs in terms of reference cell subtypes.

### Clonotype meta-analysis suggests that human tumor-specific EM CD8 T cells differentiate into exhausted/dysfunctional TILs through a progenitor subtype that upregulates *TOX*

Recent studies have shown that only a fraction of tumor-resident CD8 T cells are able to recognize tumor antigens^46,47^ and identification of tumor-reactive T cells is far from trivial. Persistent antigenic stimulation of T cells in cancer and chronic infection induce a (TOX-driven) exhaustion program that sustains high expression of inhibitory receptors. Indeed, multiple surface markers associated with exhaustion have been proposed as markers of tumor reactivity, including PD-1^45^, TIM-3^48^ and CD39^46^.

To test the assumption that human Tex cells are indeed tumor reactive, we first analyzed TIL subtype composition across tissues in the Li *et al.* melanoma cohort. As expected, the most abundant T cell subset in blood was the naïve-like (likely including naïve and central memory cells), followed by EM cells (Figure 6 A). Compared to blood, metastatic lymph node (mLN) biopsies were enriched in Tfh, Tregs, Tex and Tpex, and tumor biopsies were strongly enriched in Tex and Tpex compared to mLN and blood (Figure 6 B). Moreover, we observed that the top expanded clonotypes were mostly occupying the Tex and Tpex subtypes (Figure 6 C). The positive correlation of Tex and Tpex subtype frequency with tumor burden, as well as the enrichment of clonally expanded T cells in these compartments, is consistent with the notion that, in cancer patients, tumor-reactive TILs are mostly found in (*PDCD1*-high *TOX*-*high*) exhausted states, similarly to mouse^25,28^.

**Figure 6:**
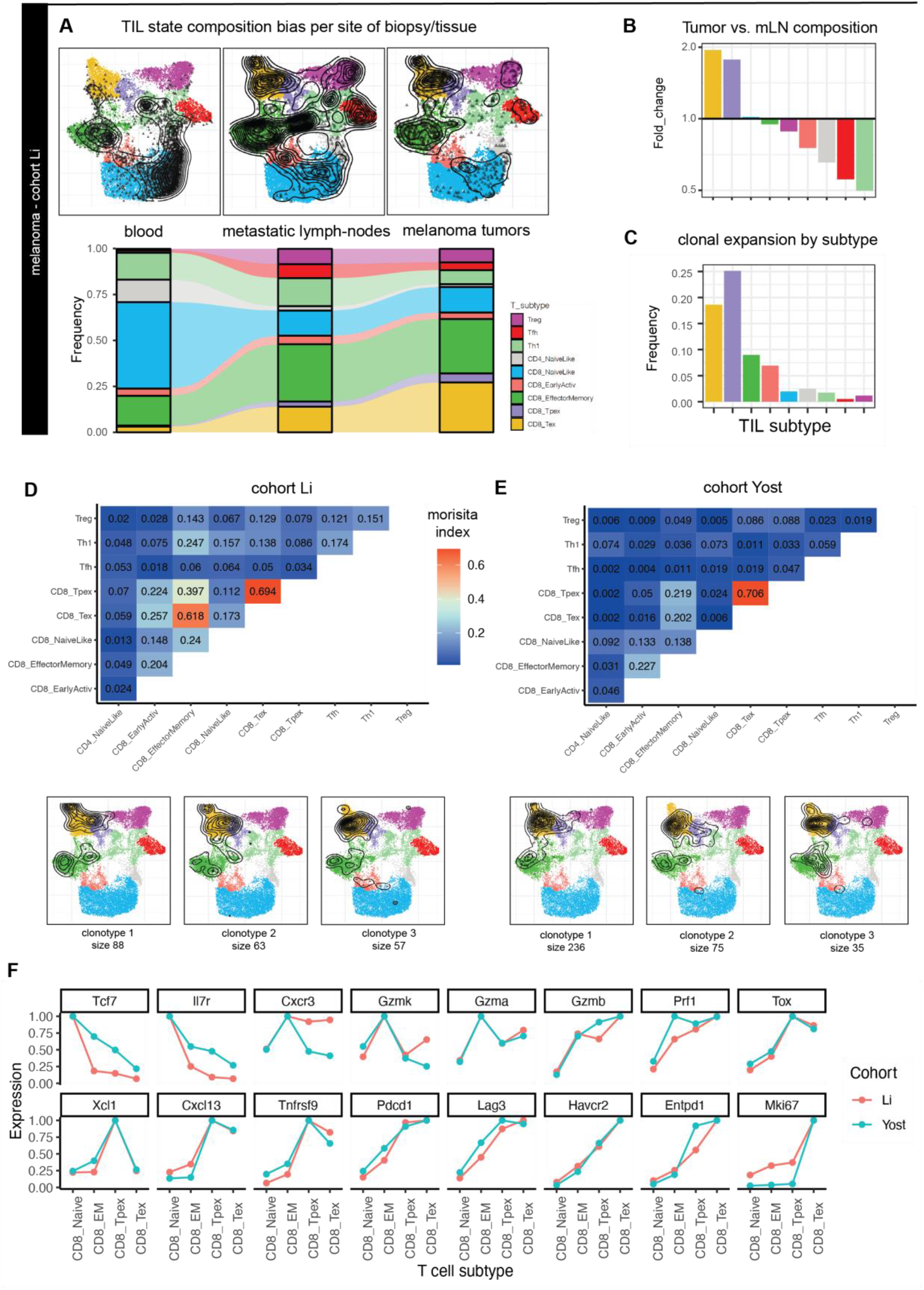
ProjecTILs analysis of human TIL states across tissues and their clonal relatedness. **A)** ProjecTILs projections and predicted subtype frequencies in biopsies from different tissues: blood, metastatic lymph nodes (mLN) and tumors (data from Li *et al.* cohort). **B)** Subtype composition bias (fold change) in tumors vs mLN. **C)** Cell frequency in each subtype occupied by the top 10 expanded clonotypes. **D-E)** The upper panels (heatmaps) display Morisita similarity indices measuring TCR repertoire overlap for each pair of TIL subtypes in the Li *et al.* (D) and Yost *et al.* (E) cohorts. Bottom panels: projection of TIL clones for the top three expanded clonotypes enriched in Tex or Tpex subtypes in each patient cohort. **F)** Average normalized gene expression of human T cells projected in the CD8_NaiveLike, CD8_EM, CD8_Tpex and CD8_Tex subtypes for a panel of key marker genes.

We next exploited TCR clonal linkage to evaluate the presence of tumor-specific TILs that do not correspond to Tex and Tpex subtypes. First, we calculated TCR clonal repertoire overlap between TIL subtypes. We verified a strong overlap between Tex and Tpex repertoires in both cohorts, as measured by Morisita similarity index (Figure 6 D-E, top panels), consistent with mice studies showing that Tpex cells give rise to Tex^17,25,28,49^. Interestingly, we also found a similarly strong clonal relatedness between Tex/Tpex and EM subtypes in the Li cohort as well as, to a lesser extent, in the Yost cohort (Figure 6 D-E). This suggested that a fraction of EM TILs were also tumor specific. Next, we selected all clonotypes that were enriched (at least 50% of the clones) in Tex or Tpex – i.e. tumor-specific clonotypes. Projection of these tumor-specific clonotypes confirmed that they spanned the three Tex, Tpex and EM subtypes in the two cohorts (Figure 6 D-E, bottom panels).

Gene expression profiles for human T cells projected onto the reference murine atlas confirmed that most key T cell markers were consistent with their ProjecTILs subtype assignment (Figure 6 F). *TOX* and *TNFRSF9 (*4-1BB) expression values were higher in Tpex and Tex compared to EM, indicating higher exhaustion and activation levels in Tpex and Tex. Consistently, *PDCD1*, *LAG3*, *HAVCR2* (TIM-3) and *ENTPD1* (CD39) were also higher in Tex compared to EM, and lower in EM compared to the naïve-like state. In contrast, *CXCR3, GZMK* and *GZMA* expression was highest in EM. Compared to Tex, tumor-specific Tpex cells expressed higher levels of *TCF7* and *IL7R*, and lower levels of cytotoxicity molecules including *GZMA, GZMB* and *PRF1*. Notably, expression of the type 1 classical dendritic cells (cDC1) chemoattractant XCL1 was specific to the Tpex subtype, consistent with Tpex gene profiling in mice^17,25,28^ and their co-localization with professional antigen-presenting cells niches in murine and human tumors^28,50^. Finally, Tpex had lower expression of cell cycling genes such as *MKI67* compared to Tex, consistent with their higher quiescence.

Altogether, these observations demonstrate that, in different human cancer types, tumor-specific CD8 TILs co-exist in three distinct subtypes: a cytotoxic *TOX*-high exhausted subtype; its *TOX*-high *TCF7*-high exhausted precursor, quiescent and characterized by lower cytotoxicity; and a precursor subtype that does not display the hallmarks of tumor-reactive TILs but resembles blood-circulating effector memory T cells. These results are compatible with a model in which *CXCR3*-high blood-circulating EM cells are recruited in the tumor, irrespectively of their antigen specificity. Then, rare tumor-reactive *TOX*-low EM TILs driven by persistent antigenic stimulation differentiate into *TOX*-high *XCL1*-high quiescent (Tpex) cells which, following interaction with XCR1+ APCs, give rise to highly proliferative terminally exhausted/dysfunctional (Tex) CD8 TILs that engage in tumor cell killing (Figure 7). Alternatively, TCR clonal linkage is compatible with a model in which some EM TILs might directly differentiate into Tex cells, without transitioning through the Tpex subtype. Importantly, our results demonstrate that these subtypes are conserved across cohorts, cancer types and species.

**Figure 7:**
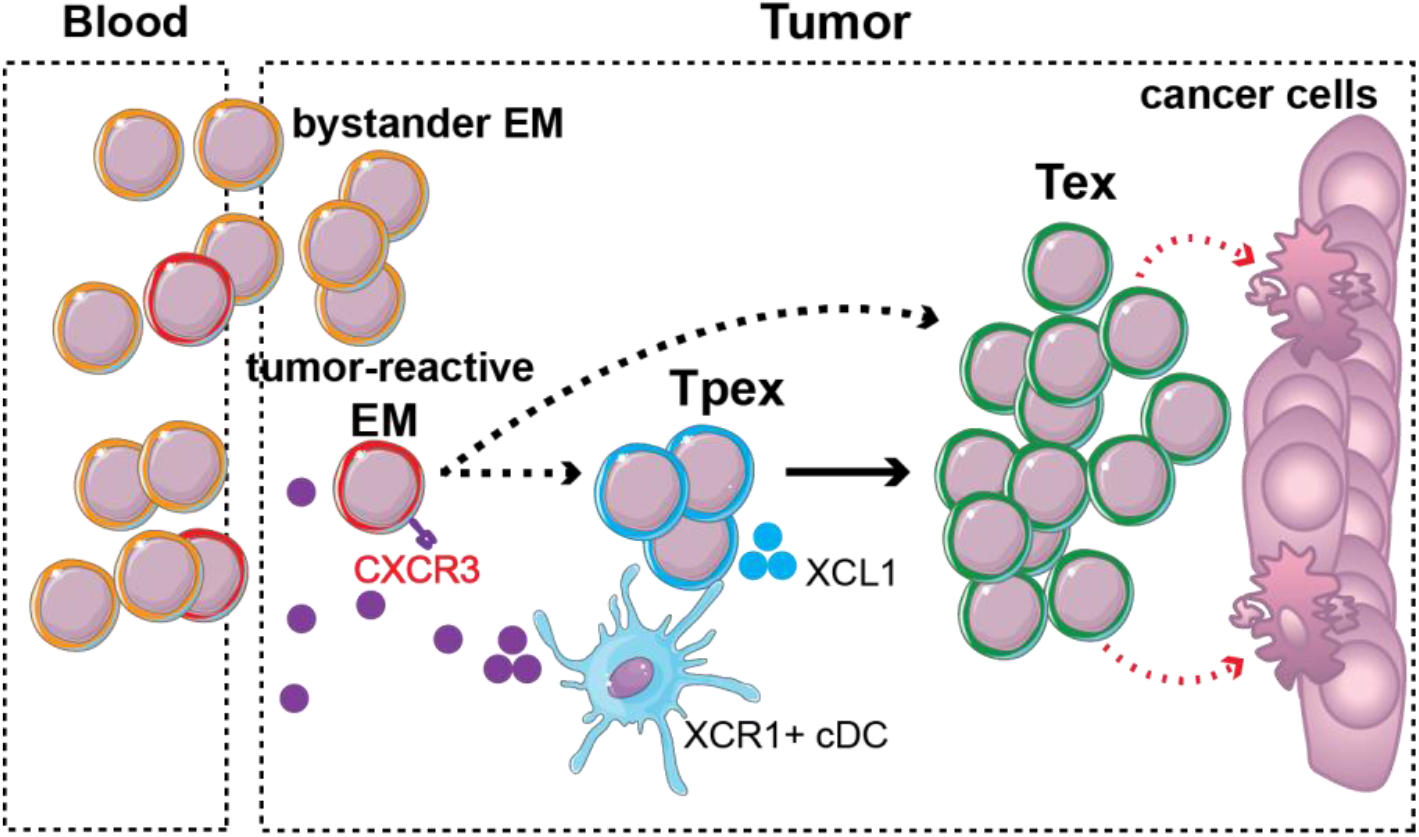
A model of intratumoral CD8 T cell differentiation supported by meta-analysis of human scRNA-seq data using ProjecTILs. Blood-circulating CXCR3-high EM cells are recruited to the tumor; these include tumor-reactive EM cells as well as bystander TILs. Persistent antigen stimulation drives differentiation of tumor-reactive EM TILs into XCL1-high Tpex cells which, following interaction with XCR1+ APCs, give rise to highly proliferative Tex CD8 TILs with capacity to kill cancer cells. An alternative differentiation path from EM directly to Tex is also plausible. **EM:** effector memory (*TOX*-low *GZMK*-high *CXCR3*-high). **Tpex**: precursor-exhausted (*TOX*-high *TCF7*-high *GZMB*-low). **Tex**: exhausted/dysfunctional (*TOX*-high, *TCF7*-low, *GZMB*-high).

## Discussion

We share the goal of many others to be able to "read" immunological states in health and disease by single-cell transcriptomics and identify therapeutic opportunities. Definition of robust, biologically relevant cellular states by scRNA-seq analysis is typically an iterative, time-consuming process that requires advanced bioinformatics and biological domain expertise. Even after successful analysis, cell clusters are not directly comparable between studies, preventing us from learning general biological rules across cohorts, conditions and models. Construction of single-cell transcriptomic atlases is a very effective approach to condense the diversity of molecular profiles within a cell type, a tissue, or an entire organism.

This is particularly powerful to characterize the heterogeneity of cell populations, to elucidate mechanisms and trajectories of differentiation as well as to aid the design of therapies targeted to specific cell types. Reference atlases can also serve as a reliable, stable baseline for the interpretation of new experiments, against which to evaluate the effect of cellular perturbations, such as changes in the balance between pre-existing subtypes or the identification of novel states in response to immunotherapies. In this work we described a novel computational method to interpret immunological states by projecting scRNA-seq data onto a reference T cell atlas, allowing the analysis of new data in the context of a stable, curated collection of T cell subtypes. This approach allows mapping T cell states that were defined across different studies, cohorts and cancer types, and provides a framework for large scale meta-analyses to identify cell states associated with prognosis and responsiveness to immunotherapy.

While it is useful to summarize cell heterogeneity as discrete subtypes for conceptualization and the design of experimental validations, it is also apparent that cells exist in a continuum of states, which would be best described as probability distributions, or regions in a multi-dimensional transcriptional space. A key advantage of embedding new data into a reference atlas of cell states is that the query cells can be interpreted in a continuous space of transcriptional states, allowing the visualization of dynamic changes between experimental conditions. While ProjecTILs can be thought of as a classifier into pre-annotated discrete reference states, it also operates in a continuous space and can, therefore, capture intermediate and transient cellular states. For instance, we observed a pattern of expression of the chemokine receptor *Cx3cr1* that straddles the Tpex and Tex subtypes in the TIL atlas (Figure 1 F), as well as a gradient of expression for this molecule within the Tex compartment in infection. Indeed, recent studies have identified *Cx3cr1* as a marker for a transitory, intermediate state between the Tpex and Tex populations^51–53^.

Previously developed methods such as *scmap*^54^, based on nearest neighbor search, allow ‘matching’ query cells to annotated cells or clusters that are part of a reference dataset, enabling the classification of new data into predefined cell types. Beyond the interpretation of new data in terms of known, annotated cell subtypes, we have shown that ProjecTILs can aid the discovery of novel states that deviate from the reference, *e.g.* as a consequence of a genetic alteration. A case in point, analysis of the *Regnase-1* KO data of Wei and colleagues not only explained the increased T cell persistence and tumor control in terms of changes in T cell subtypes frequencies (Figure 3 E) but also revealed the previously unreported down-regulation of a novel inhibitory gene program in exhausted CD8 TILs. This program (ICA 25) was driven by *Klrc1*/NKG2A and *Lag3* (Figure 3 G-H), and its expression was uncoupled from that of other inhibitory receptors such as PD-1, TIM-3 or CTLA4, suggesting a potential benefit of targeting these two programs simultaneously. Indeed, dual blockade of PD-1 and LAG-3 results in robust and synergistic reinvigoration of Tex cells in cancer^55^ and chronic infection^56^, while NKG2A blockade has been recently shown to potentiate CD8 T cell immunity induced by cancer vaccines^57^.

In most experiments aimed at studying the effect of a perturbation, it is imperative to design a control group, as a baseline to compare the effect of the perturbation of interest. In single-cell experiments, when studying the effect of a genetic perturbation or of a treatment, the control group usually consists of the same cell population as the perturbation group but under basal conditions. It is conceivable, as reference atlases become increasingly complete, that the control group is already satisfactorily included in the reference atlas – the new condition could then be directly evaluated against the atlas, bypassing the need for deeply re-sampling the transcriptional space of basal conditions, as we have illustrated with the analysis of Regnase-1 KO data in absence of a control sample (Suppl. Figure S8).

Compared to murine models, the analysis of human TIL states is complicated by the large genetic and environmental variability between patients, as well as the large variability between biopsies due to tissue-specific effects and tumor heterogeneity. Tumor scRNA-seq studies typically describe T cell heterogeneity in terms of several clusters, which are then manually annotated in “states” or “subtypes” defined by the authors and that tend to suffer from batch effects between samples. As a result, systematic comparison of T cells states across studies, cohorts and cancer types becomes extremely difficult. In this work, we have shown that ProjecTILs can accurately project human T cell transcriptomes onto a reference mouse atlas, and that human TIL heterogeneity can be largely explained in terms of robust T cell subtypes. Such remarkable level of conservation between human and mouse TIL states is extremely encouraging for translational research in cancer immunotherapy.

Preventing or reverting exhaustion/dysfunction of tumor-specific CD8 T cells is currently one of the major goals in cancer immunotherapies. While there is evidence suggesting that pre-exhausted/dysfunctional tumor-specific CD8 T cells are present in human tumors, a robust definition of such TIL states and the differentiation process by which they acquire exhaustion features has remained elusive. Our meta-analysis of scRNA-seq and TCR-seq data from two cohorts of melanoma and basal cell carcinoma patients with ProjecTILs, revealed that i) the majority of human TILs do not display features of exhaustion or tumor-reactivity, and are clonally disconnected from the exhausted TILs, suggesting that most of them are not tumor-reactive; and that ii) tumor-specific exhausted/dysfunctional CD8 TILs can co-exist with two rare, quiescent precursor subtypes: an exhausted TOX+ PD1+ TIM3− XCL1+ (Tpex) state; and a pre-exhausted/dysfunctional (EM) state with low expression of *TOX* and inhibitory receptors, and high expression of *CXCR3* and *GZMK,* that resemble blood circulating EM cells. The CD8 TIL differentiation model proposed based on these observations has important implications for the design of therapies aimed at preventing T cell exhaustion, and for the identification of tumor-specific T cells with high stemness for their use in adoptive cell therapies^58^.

We have described the construction of reference single-cell atlases for murine T cells in pan-cancer and infection models that are strongly supported by literature. While we observed that the main, known T cell subtypes can be accurately recapitulated in these reference maps, they do not yet encompass the full diversity of transcriptional states that can be acquired by T cells, especially for CD4 TILs (which were under-represented compared to CD8 among available data) and for γδ T cells (which were not represented at all). Only very recently, with the popularization of single-cell technologies, it has become possible to generate data of sufficient depth and quality to construct such high-resolution reference maps. We are therefore just beginning to appreciate the full potential of combining information from multiple studies and perform meta-analyses across models, tissues and cancer types. Considering the pace at which new single-cell data are generated, we anticipate that reference maps will quickly grow in size and completeness, increasingly covering the space of possible transcriptional states that can be assumed by individual cells. We expect that the accuracy of projection of new data into such exhaustive reference atlases will also improve as a consequence.

While we have shown that ortholog mapping offers a viable solution to interpret human T cell responses in the context of a robust mouse atlas, we envision that, with rapidly growing data, it will soon become feasible to construct high-quality reference human T cell atlases able to capture human-specific diversity. In this respect, mouse atlases could serve as scaffolds to build their human counterparts. Finally, projecting whole tissue data into a collection of high-resolution cell type atlases, covering multiple immune cell compartments, would enable interpreting immune responses at a systems level, by the study of correlation, and putative interaction and cross-talk, between not only cell types, but between cells in very specific differentiation states.

We have implemented ProjecTILs as an R package (https://github.com/carmonalab/ProjecTILs) and we provide a Docker image ready to use. Because ProjecTILs is integrated with Seurat 3, it can be easily combined with other tools for up- and down-stream analyses. We believe our approach will have a great impact in revealing the mechanisms of action of experimental immunotherapies and to guide novel therapeutic interventions in cancer and beyond.

## Methods

### Single-cell RNA-seq of tumor-draining lymph node T cells

Eight-to 10-week-old female C57Bl/6 mice were obtained from The Charles River Laboratories and housed at Genentech in standard rodent micro-isolator cages to be acclimated to study conditions for at least 3 days before tumor cell implantation. Tumor draining lymph nodes from mice with established MC38 tumors (~190 mm^3^) were excised and single-cell suspensions were stained for CD45 (Biolegend, clone 30-F11, dilution 1:100), TCRb (Biolegend, clone H57-597, 1:100), CD44 (eBioscience, clones IM7, 1:200), CD62L (eBioscience clone Mel14, 1:80) and LIVE/DEAD (Life Technologies, Fixable Dead Cell Stain) and sorted into CD45^+^TCRb^+^ T cells, gating out the antigen-inexperienced CD62L+CD44-population. Sorted cells were then loaded onto a 10x Chromium Chip A using reagents from the Chromium Single-Cell 5’ Library and Gel Bead Kit (10x Genomics) according to the manufacturer’s protocol. Amplified cDNA was used for both 5’ RNA-seq library generation and TCR V(D)J targeted enrichment using the Chromium Single-Cell V(D)J Enrichment Kit for Mouse T Cells (10x Genomics). 5’ RNA-seq and TCR V(D)J libraries were prepared following the manufacturer’s user guide (10x Genomics). The final libraries were profiled using the Bioanalyzer High Sensitivity DNA Kit (Agilent Technologies) and quantified using the Kapa Library Quantification Kit (Kapa Biosystems). Each single-cell RNA-seq library was sequenced in one lane of HiSeq4000 (Illumina) to obtain a minimum of 20,000 paired-end reads (26 × 98 bp) per cell. Single-cell TCR V (D)J libraries were multiplexed and sequenced in one lane of HiSeq2500 (Illumina) to obtain minimum of 5,000 paired-end reads (150 × 150 bp) per cell. The sequencing specifications for both single-cell RNA-seq and TCR V(D)J libraries were according to the manufacturer’s specification (10x Genomics).

Single-cell RNA-seq data for each replicate were processed using cellranger *count* [CellRanger 2.2.0 (10x Genomics)] using a custom reference package based on mouse reference genome GRCm38 and GENCODE^59^ gene models. Individual count tables were merged using CellRanger *aggr* to reduce batch effects.

### Cell lines

The murine colon adenocarcinoma MC38 cell line was obtained from a former Genentech colleague, Rink Offringa, in 2008. Cells were cultured in RPMI-1640 medium plus 2 mmol/L l-glutamine with 10% fetal bovine serum (HyClone). Cells in log-phase growth were centrifuged, washed once with Hank’s balanced salt solution (HBSS), counted, and resuspended in 50% HBSS and 50% Matrigel (BD Biosciences) at 1 × 10^6^ cells/mL for injection into mice.

### Public Single-cell RNA-seq data

To construct the reference TIL atlas, we obtained single-cell gene expression matrices from the following GEO entries: GSE124691^21^, GSE116390^17^, GSE121478^31^, GSE86028^60^; and entry E-MTAB-7919^23^ from Array-Express. Data from GSE124691 contained samples from tumor and from tumor-draining lymph nodes, and were therefore treated as two separate datasets. For the TIL projection examples (OVA Tet+, miR-155 KO and Regnase-KO), we obtained the gene expression counts from entries GSE122713^25^, GSE121478^31^ and GSE137015^32^, respectively.

Single-cell data to build the LCMV-specific CD8 T cell reference map were downloaded from GEO under the following entries: GSE131535^35^, GSE134139^36^ and GSE119943^37^, selecting only samples in wild type conditions. Data for the Ptpn2-KO, Tox-KO and CD4-depletion projections were obtained from entries GSE134139^36^, GSE119943^37^, and GSE137007^42^ and were not included in the construction of the reference map.

Single-cell human data for ortholog projection was downloaded from GEO under the following entries: GSE123139^5^, GSE123813^44^ and GSE120575^6^.

### Batch-effect correction and construction of reference atlases

Prior to dataset integration, single-cell data from individual studies were filtered using TILPRED-1.0 (https://github.com/carmonalab/TILPRED), which removes cells not enriched in T cell markers (e.g. *Cd2*, *Cd3d*, *Cd3e*, *Cd3g*, *Cd4*, *Cd8a*, *Cd8b1*) and cells enriched in non-T cell genes (e.g. *Spi1*, *Fcer1g*, *Csf1r*, *Cd19*). Dataset integration was performed using STACAS^14^ (https://github.com/carmonalab/STACAS), a batch-correction algorithm based on Seurat 3^12^. For the TIL reference map, we specified 600 variable genes per dataset, excluding cell cycling genes, mitochondrial, ribosomal and non-coding genes, as well as genes expressed in less than 0.1% or more than 90% of the cells of a given dataset. For integration, a total of 800 variable genes were derived as the intersection of the 600 variable genes of individual datasets, prioritizing genes found in multiple datasets and, in case of draws, those derived from the largest datasets. We determined pairwise dataset anchors using STACAS with default parameters, and filtered anchors using an anchor score threshold of 0.8. Integration was performed using the *IntegrateData* function in Seurat3, providing the anchor set determined by STACAS, and a custom integration tree to initiate alignment from the largest and most heterogeneous datasets. Similarly, to construct the LCMV reference map, we split the datasets into five batches that displayed strong technical differences, and applied STACAS to mitigate their confounding effects. We computed 800 variable genes per batch, excluding cell cycling genes, ribosomal and mitochondrial genes, and computed pairwise anchors using 200 integration genes, and otherwise default STACAS parameters. Anchors were filtered at the default threshold 0.8 percentile, and integration was performed with the *IntegrateData* Seurat3 function with the guide tree suggested by STACAS.

Both for the TIL and LCMV atlases, we performed unsupervised clustering of the integrated cell embeddings using the Shared Nearest Neighbor (SNN) clustering method^61^ implemented in Seurat 3 with parameters {*resolution*=0.6, *reduction*=”umap”, *k.param*=20} for the TIL atlas and {*resolution*=0.4, *reduction*=”pca”, *k.param*=20} for the LCMV atlas. We then manually annotated individual clusters (merging clusters when necessary) based on several criteria: i) average expression of key marker genes in individual clusters; ii) gradients of gene expression over the UMAP representation of the reference map; iii) gene-set enrichment analysis to determine over- and under-expressed genes per cluster using MAST^62^. In order to have access to predictive methods for UMAP, we recomputed PCA and UMAP embeddings independently of Seurat3 using respectively the *prcomp* function from basic R package “stats”, and the “umap” R package (https://github.com/tkonopka/umap).

### The ProjecTILs pipeline

The essential input to the ProjecTILs pipeline is an expression matrix, where genes are rows and cells are columns. If raw counts (e.g. UMI counts) are provided, each entry *x* in the matrix will be normalized using the formula: log (1 + 10,000 x / S), where S is the sum of all counts for that cell, and log is the natural logarithm. To ensure that only T cells are included in the query dataset, by default TILPRED-1.0 is applied to predict the composition of the query, and all cells annotated as “Non-T cells” or “unknown” are removed from the query. This filter can be optionally disabled by the user. Then, a reference atlas of annotated cells states (by default the TIL atlas) is loaded into memory, together with its cell embeddings in gene, PCA and UMAP spaces, and all associated metadata. In order to bring the query data in the same representation spaces as the reference map, batch-effect correction is applied to the normalized cell-gene counts of the query set using the anchor-finding and integration algorithms implemented in STACAS and Seurat3, where the genes for integration consist of the intersection of the variable genes of the reference map and all genes from the query. After batch-effect correction, the PCA rotation matrix pre-calculated on the reference atlas (i.e. the coefficients allowing the transformation from reference gene space into PCA space) is applied to the normalized, batch-corrected query matrix. In the same way, the *predict* function of the “umap” package allows transforming PCA embeddings into UMAP coordinates. By these means, the query data can be embedded into the original, unaltered coordinate spaces of the reference atlas, enabling joint visualization as well as classification of the query cells into T cell subtypes. ProjecTILs is implemented as a modular R package, with several functions that aid interpretation and analysis. The *make.projection* function is the core utility that implements the projection algorithm described above. It can be run in “direct” mode, in which case the PCA and UMAP rotations are directly applied without batch-effect correction. This may be useful for very small datasets, where alignment and integration algorithms will not be applicable. To project human data onto a murine reference atlas, the user must set the flag “human.ortho=TRUE”, which automatically converts human genes to their mouse orthologs before projection. *Plot.projection* allows visualizing the query dataset as density level curves superimposed on the reference atlas. The *cellstate.predict* function implements a nearest-neighbor classifier, which predicts the state of each query cell by a majority vote of its annotated nearest neighbors (either in PCA or UMAP space) in the reference map. *Find.discriminant.dimensions* analyses PCA and ICA embeddings (described below) to determine dimensions where the query deviates significantly from the reference map. Several additional functions allow visualizing multiple aspects of the reference and projected dataset and aid the biological interpretation of the results. The code and description of the package, together with tutorials and applications to analyze public datasets can be found at: https://github.com/carmonalab/ProjecTILs

### ICA and discriminant dimensions

Independent component analysis (ICA) is a computational technique aimed at deconvoluting a multivariate signal (such as simultaneous expression of many genes) into additive, independent sources (in this case different genetic programs). We applied the fastICA implementation^63^ to determine 50 independent components on the integrated expression matrix of the TIL reference atlas. To suggest a biological interpretation of the ICA components, we downloaded hallmark gene sets (H) and canonical pathway gene sets (CP) from the Molecular Signatures Database^34^ (mSigDB), as well as selected immunological signatures from previous studies. We scored each ICA against these signatures, retaining the top three-scoring signatures for each ICA, and clustered ICA components based on the union of all retained signatures (see Suppl. Figure S5).

After projection, query datasets are also subject to transformation in ICA space through the ICA rotation matrix. The *find.discriminant.dimensions* function implemented in ProjecTILs evaluates the distribution of the cells in the query for each ICA dimension, and compares it to the distribution of cells in the reference (or to a control query dataset, if provided) in the same ICA dimension. For each ICA dimension, a statistical test can then be applied to confirm or reject the null hypothesis that, in this dimension, the cells in reference and query are drawn from the same distribution. ProjecTILs implements a Kolmogorov-Smirnov (KS) test or a t-test to the null hypothesis, multiplying p-values by the number of tests (i.e. 50) to correct for multiple testing (*i.e.* Bonferroni correction). ICA dimensions where the query deviates significantly from the reference (or the control, if provided) are ranked by their test statistic to determine the top discriminant dimensions. ICA embeddings can be visualized as an additional dimension on the z-axis of the reference UMAP space with the function *plot.discriminant.3d*.

### Cross-validated projection benchmark

To estimate the accuracy of the projection algorithm, we devised a cross-validation experiment where we removed part of the data from the reference before projecting the removed data back into the map. Each of the seven datasets included in the reference TIL map, except one, was composed of at least two samples; we constructed a cross-validation experiment by removing, at each cross-validation step, half of the samples of a given dataset, and then projected these samples into a reduced version of the map that does not contain the data points from these samples. The dataset by Singer *et al.*^60^ consists of a single sample, and therefore we removed all of its cells before projecting it back into its reduced map. After systematically projecting all cells in cross-validation, we compared their projected coordinates (either in UMAP or PCA space) with their original coordinates in the reference map. By the same token, we evaluated the performance of the projection algorithm by comparing the composition of any given dataset in terms of predicted cell states compared to the annotated cell states in the reference map (Suppl. Figure S2). We found that using ProjecTILs with batch-correction, 90.0% of the projected cells are found within a radius of one unit (their neighborhood) in UMAP space (Suppl. Figure S3) from their original coordinate in the reference map, and 93.4% within a radius of 5 units in PCA space. In terms of cell state classification, 91.6% of the projected cells were correctly assigned to their cell subtype (Suppl. Figure S2 A).

### Human-mouse ortholog projection

We downloaded the list of orthologous genes between human and mouse from Ensemble BioMart release 101. When mapping was ambiguous (i.e. a human gene mapping to several murine genes and viceversa) we favored the identical upper-case human ortholog translation of mouse genes, when available. When projecting human scRNA-seq data onto a mouse reference atlas with ProjecTILs (human.ortho=TRUE), the expression matrix was automatically subset on the human genes with a valid ortholog, and submitted for projection using the mouse gene identifiers. All subsequent analyses were performed in the mouse ortholog space.

Clonal overlap between subtypes was calculated using the Morisita index implementation of scRepertoire^24^. Because of the large imbalance in terms of cell subtypes in the Yost et al. cohort, we uniformly down-sampled all subtypes to a maximum of 500 cells for clonal overlap calculation. For the analysis of gene expression profiles in the Li et al. and Yost et al. cohorts, we calculated the average expression of all cells projected in the CD8_NaiveLike, CD8_EM, CD8_Tpex and CD8_Tex subtypes for 16 key marker genes. To be able to compare the profile of different genes, we rescaled the average expression value of a given gene by its maximum average expression over the four subtypes under analysis (CD8_NaiveLike, CD8_EM, CD8_Tpex and CD8_Tex).

### Animal study oversight

All animal studies were reviewed and approved by Genentech’s Institutional Animal Care and Use Committee. Mice whose tumors exceeded acceptable size limits (2,000 mm3) or became ulcerated were euthanized and removed from the study.

## Accession Codes

Generated scRNA-seq data of MC38 tumor-draining lymph node T cells (excluding antigen-inexperienced CD62L+CD44-cells) were deposited in the ArrayExpress database with accession ID E-MTAB-9274.

Published scRNA-seq data of MC38 TILs: E-MTAB-7919, GSE124691.

Published scRNA-seq data of B16 TILs GSE116390, GSE121478, GSE86028, GSE122713, GSE137015.

Published scRNA-seq data of LCMV infection P14 cells: GSE131535, GSE134139, GSE119943, GSE119943, GSE137007.

Published scRNA-seq data from cancer patients (melanoma and basal cell carcinoma): GSE123139, GSE123813, GSE120575.

## Acknowledgements

We are grateful to Dr. Amaia Martinez-Usatorre, Prof. Werner Held and Dr. Gonzalo Parra for critical reading of the manuscript. This research was possible thanks to the support of the Swiss National Science Foundation (SNF) Ambizione grant 180010 to SJC. SJC would like to thank the SNF for supporting researchers developing their early independent academic careers in Switzerland, regardless of their nationalities.

## Supplementary Material

**Supplementary Figure S1:**
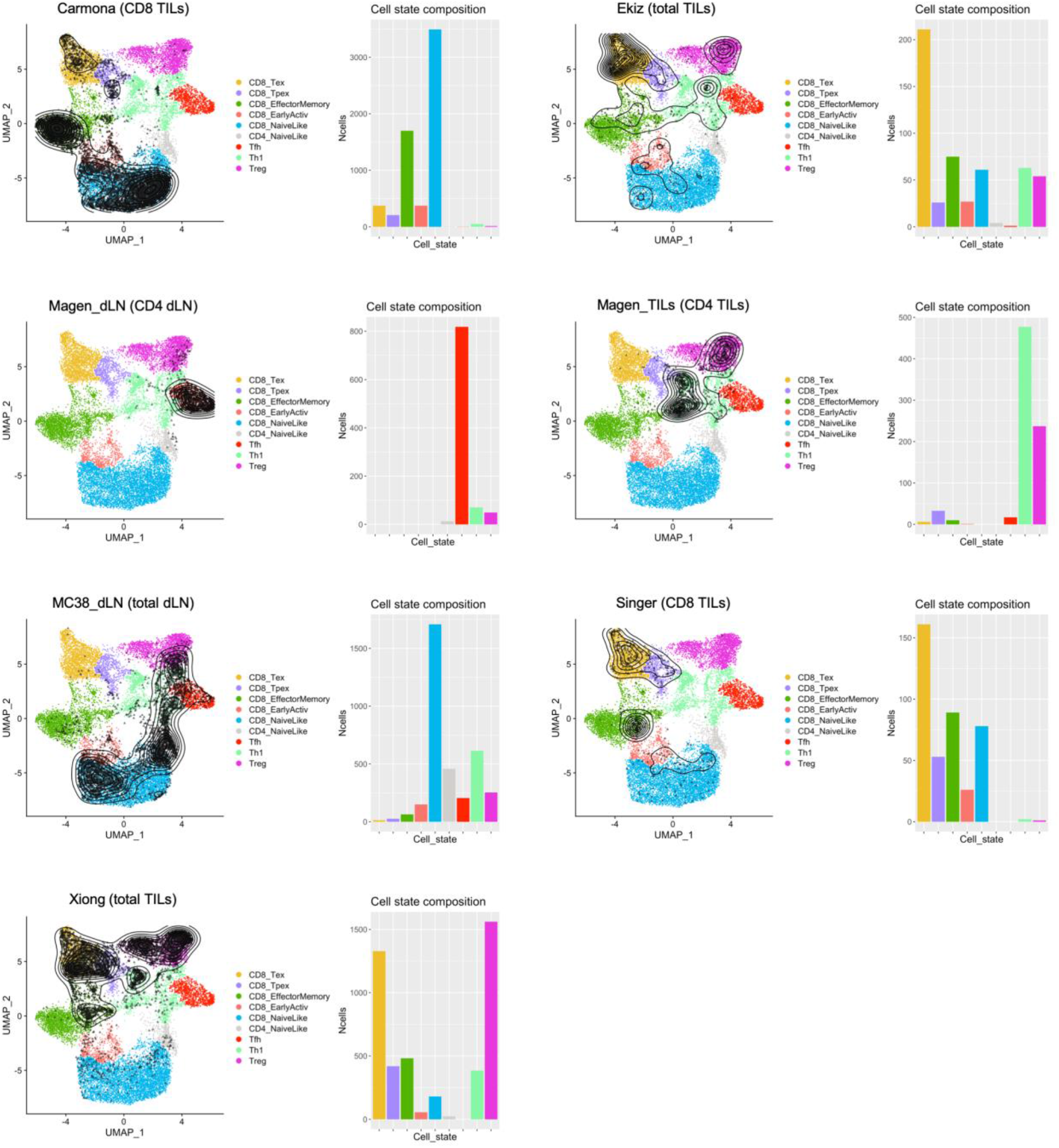
Distribution of samples on the reference TIL atlas. For each of the seven datasets composing the reference atlas, individual cells from the given dataset are plotted as black points on the reference map, and point density is summarized using level lines. Barplots quantify the number of cells included in each T cell subtype for the given dataset. TILs: tumor-infiltrating lymphocytes; dLN: draining lymphnode.

**Supplementary Figure S2:**
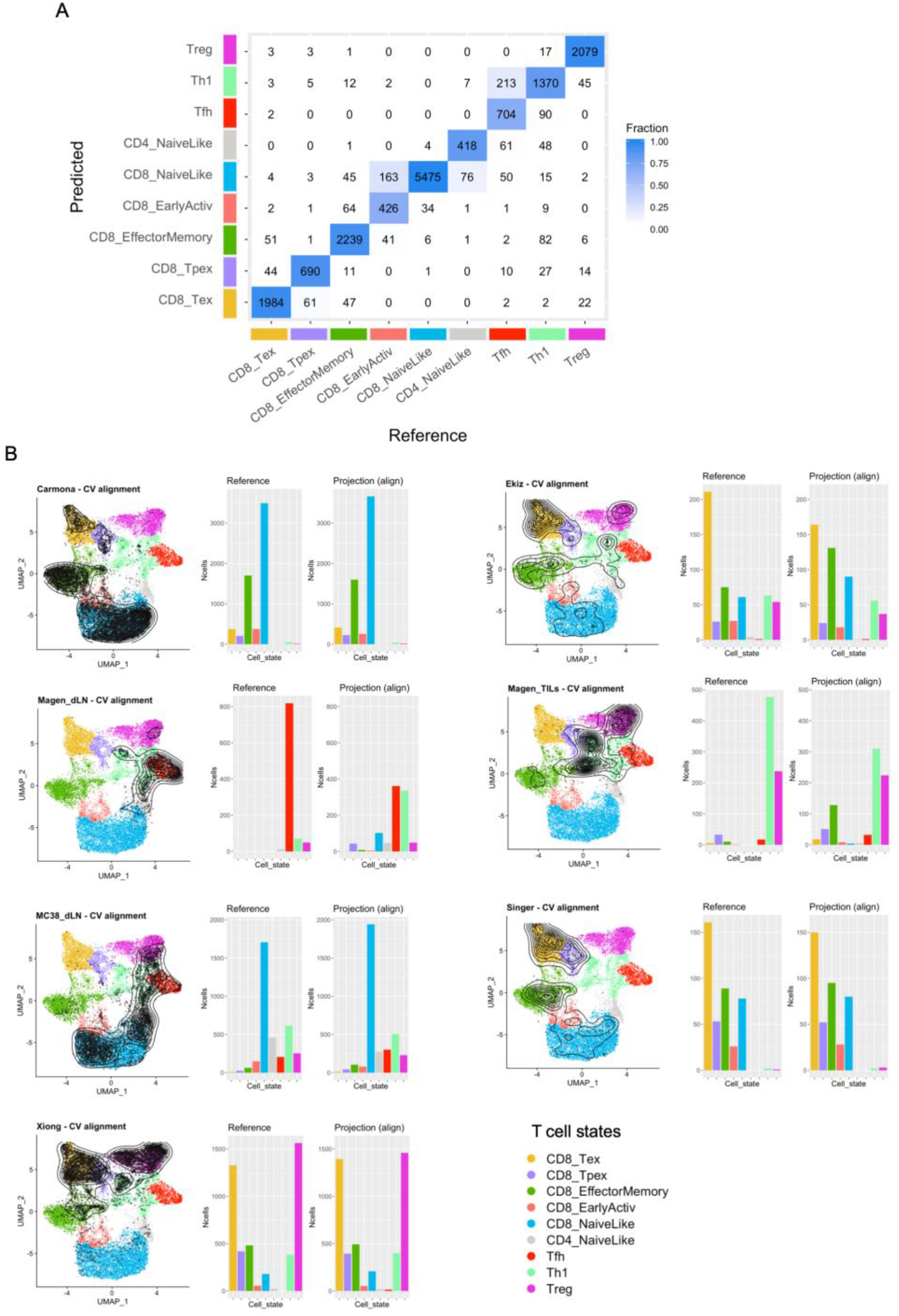
Cell state composition for reference and projection in cross-validation benchmark. **A)** Confusion matrix for ProjecTILs state classification in cross-validation, comparing reference (x-axis) vs predicted (y-axis) states. The numbers indicate the number of cells for each pair of reference/predicted states (the diagonal contains the correctly classified cells, accounting for 91.6% of the total). **B)** For each dataset, the projected coordinates in cross-validation are shown as level curves on the UMAP representation of the map (left). Based on the projected coordinates, a nearest-neighbor algorithm is applied to predict the cell state of the query cells; the predicted composition of the dataset can be then compared to the “true” composition of the reference map for the same dataset (right).

**Supplementary Figure S3:**
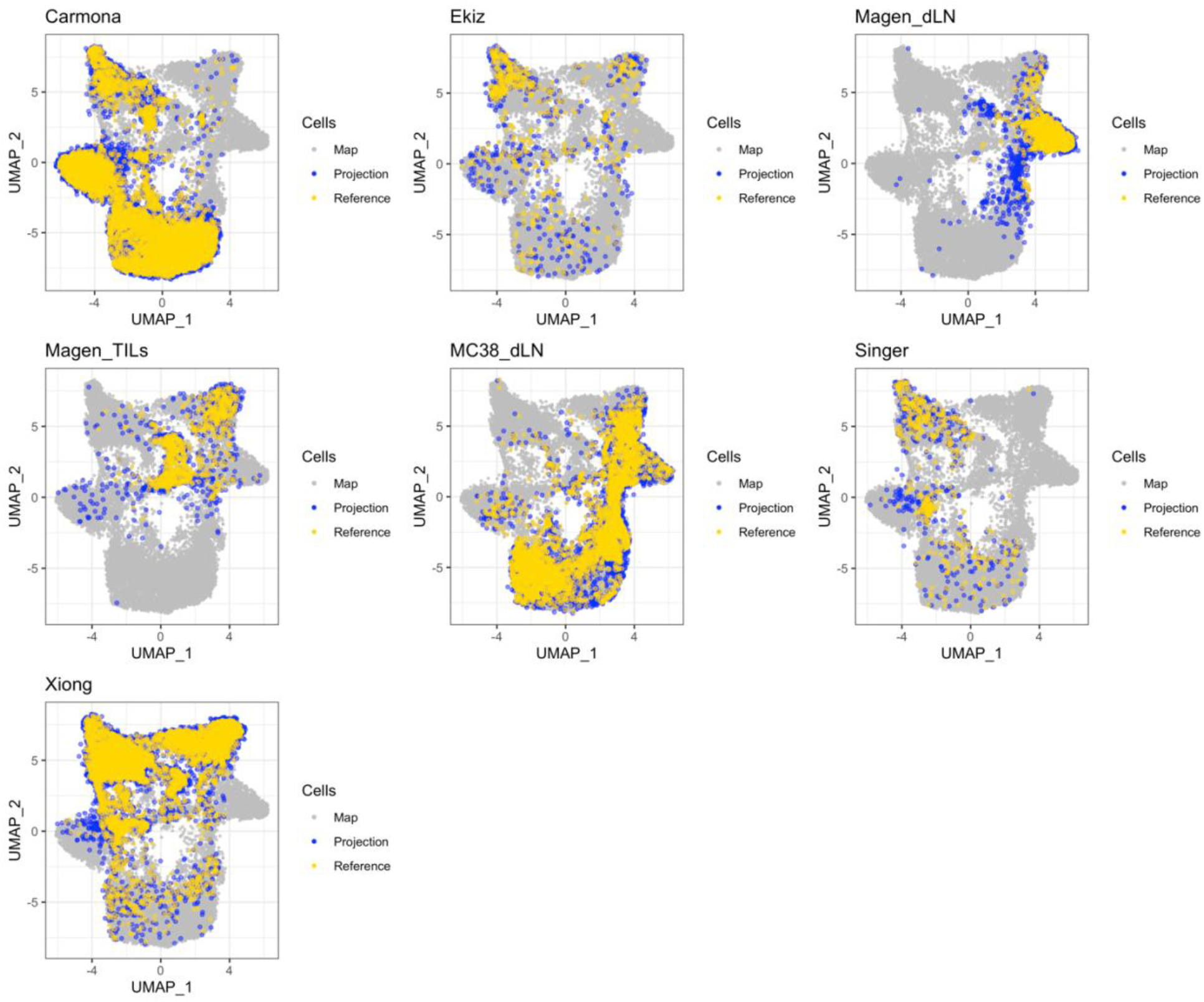
Cross-validated projection of dataset in the reference map. For each of the seven datasets included in the reference TIL map, we constructed a cross-validation experiment by removing, at each cross-validation step, half of the samples of a given dataset, and then projected these samples into a reduced version of the map that does not contain the data points from these samples. The projected coordinates (blue) can be compared against the original coordinates in the map (yellow); the complete map layout is shown in gray. Note that the distribution pattern of each dataset is associated to the specific T cell subtypes it contains, e.g. datasets Carmona and Singer contain CD8 TILs, Magen_TILs contain CD4 TILs, Ekiz and Xiong contain both CD4 and CD8 TILs, Magen_dLN and MC38_dLN contain CD4 T cells, and both CD4 and CD8 T cells, respectively, from tumor-draining lymph nodes.

**Supplementary Figure S4:**
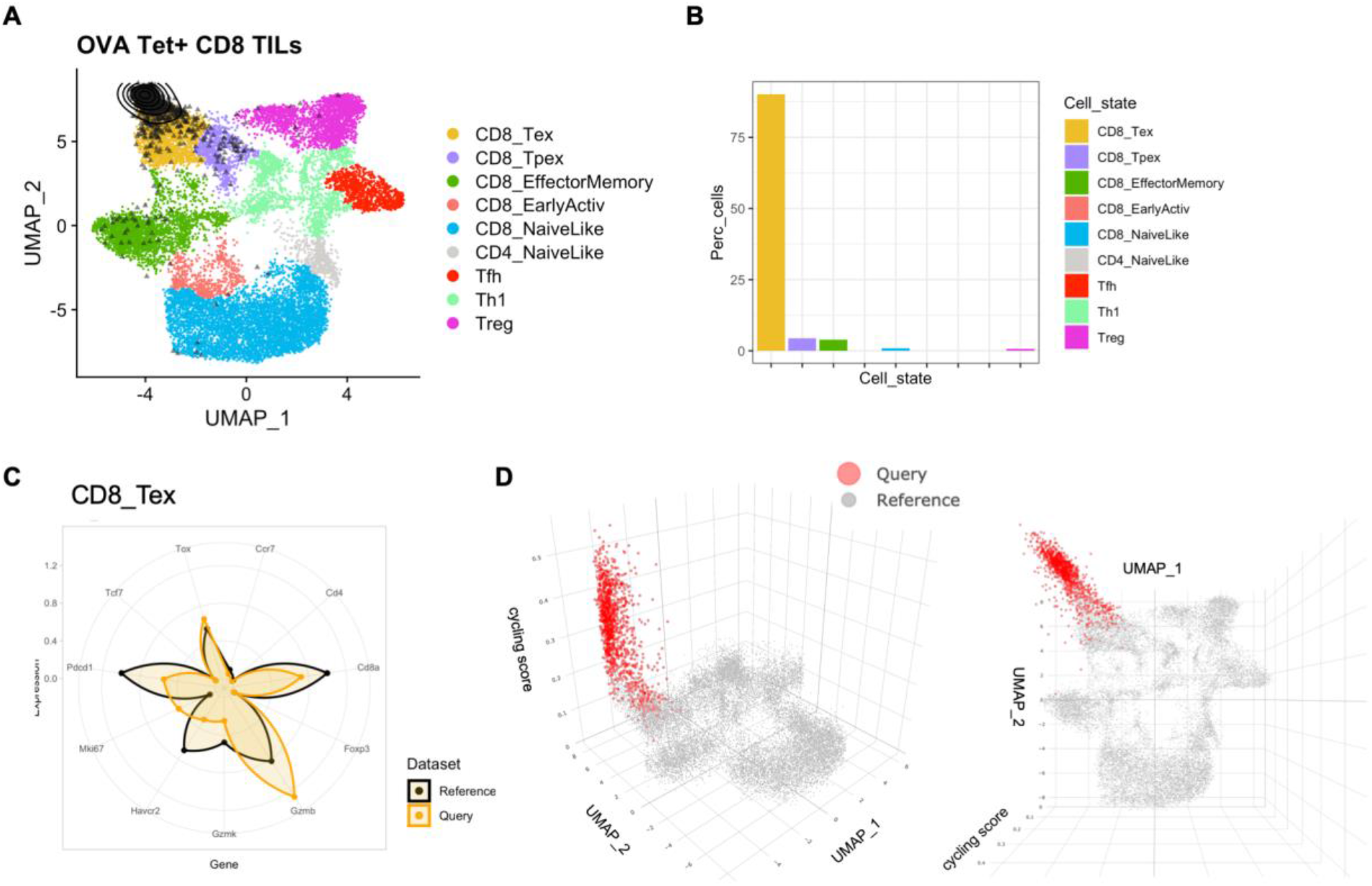
Projection of tumor-specific tetramer+ CD8 TIL single-cell data from Miller et al. **A)** Predicted coordinates in UMAP space as density contours; **B)** percentage of cells projected in each of the functional clusters of the reference atlas**; C)** gene expression signature for predicted CD8 terminally exhausted (CD8_Tex) cells; **D)** cell cycling score plotted on the z-axis of the UMAP plot (side and top view).

**Supplementary Figure S5:**
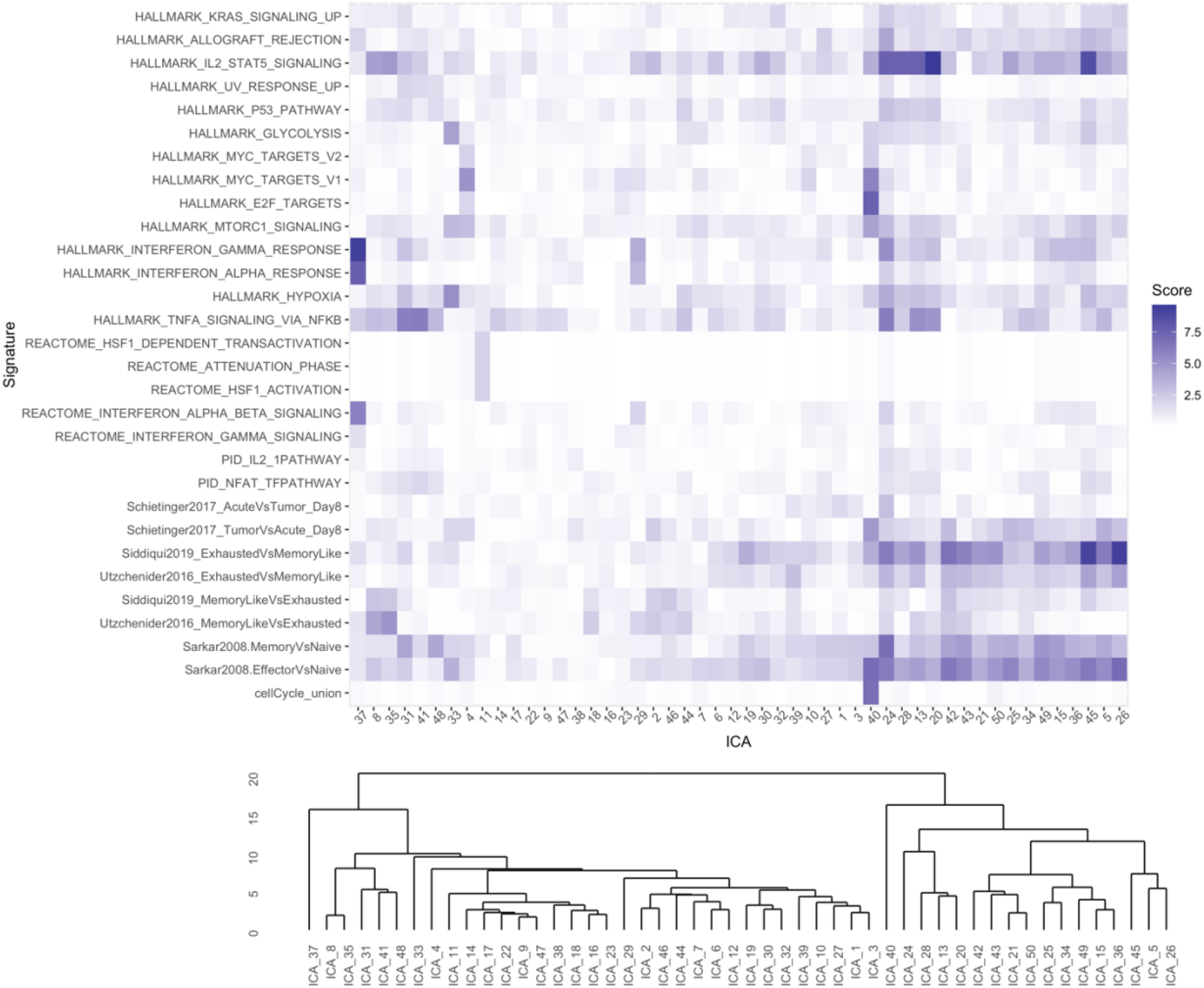
Correlation between ICA genes and annotated signatures from mSigDB. Score for each ICA component against a selection of gene signatures from the Molecular Signature Database (top). Dendrogram of ICA components clustered by signature similarity (bottom).

**Supplementary Figure S6:**
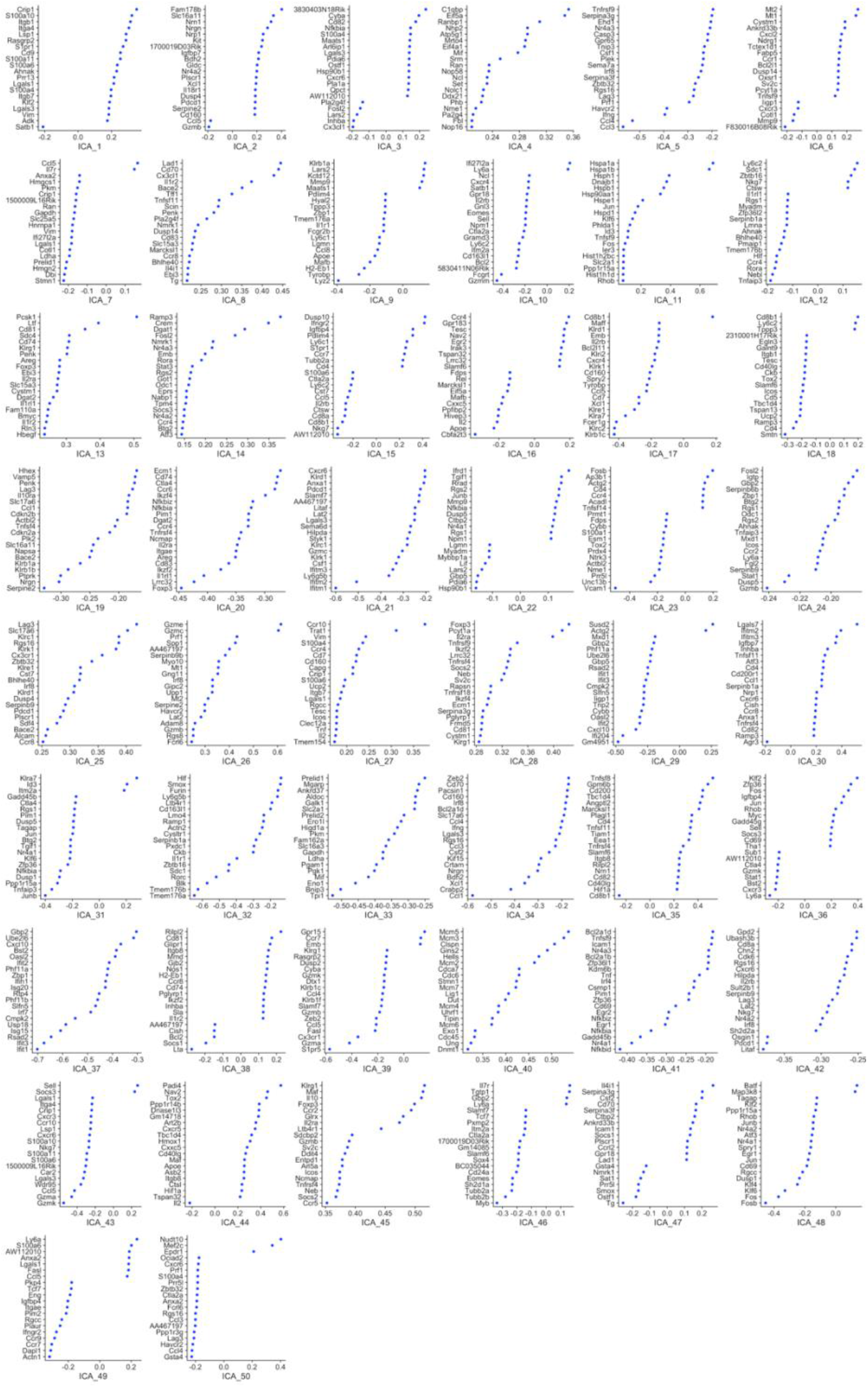
Top 20 genes associated with each of the 50 ICA dimensions of the reference TIL atlas. The scale on the x-axis represents the ICA loading of each gene, and gives an indication of the contribution of each gene to the cell coordinate in the given ICA component.

**Supplementary Figure S7:**
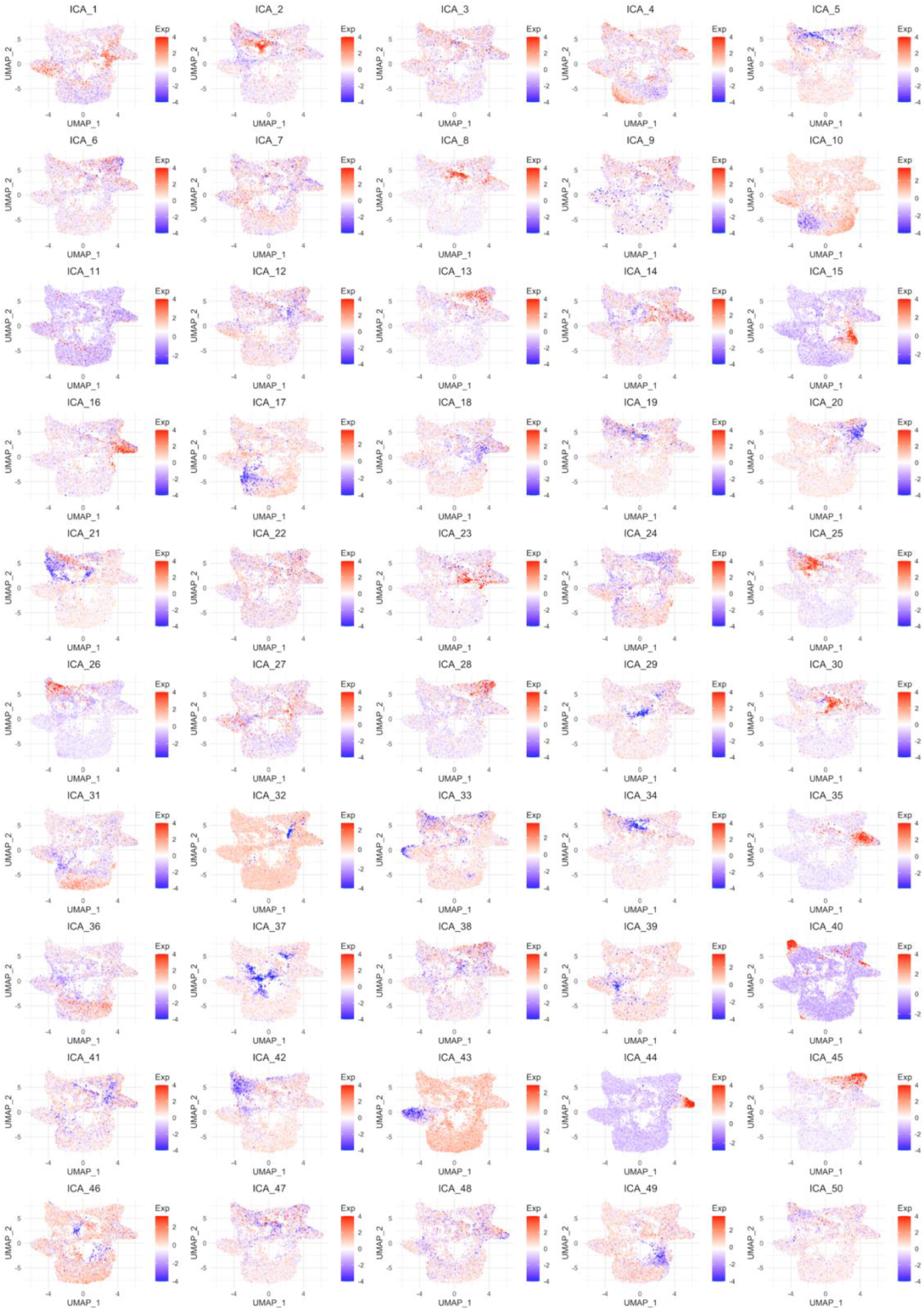
Reference cell embeddings in the 50 ICA dimensions of the reference TIL atlas. The Exp scale represents the coordinate of each cell for the given ICA component.

**Supplementary Figure S8:**
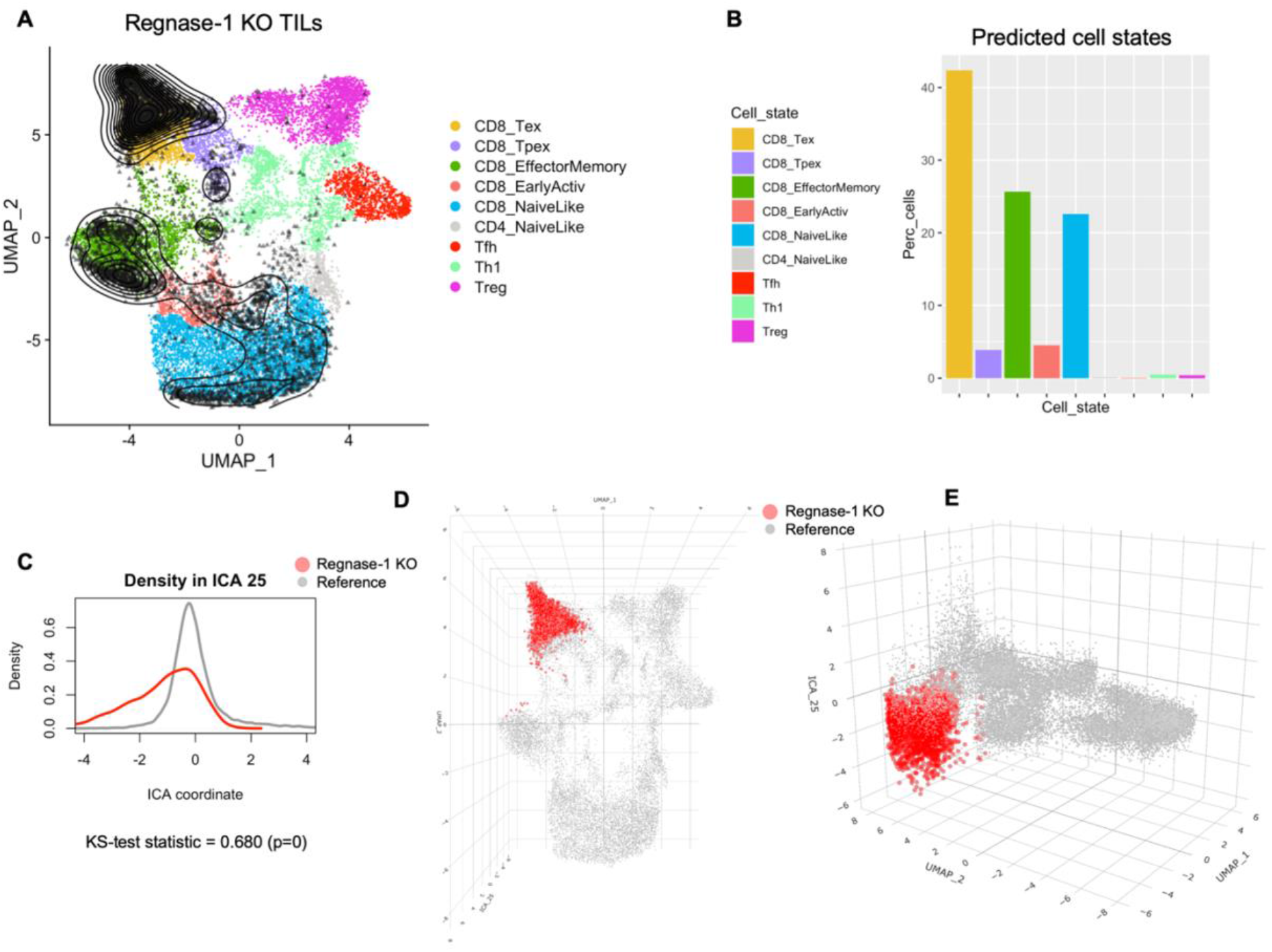
**A)** Projection of Regnase-1 KO single-cell data onto the TIL reference atlas, without control sample. **B)** Predicted cell subtype composition in terms of nearest neighbors on the reference map. **C)** ICA component 25 was ranked as the second most significant by KS-test (statistic=0.680, p=0) for the null hypothesis that ICA 25 embeddings for CD8 exhausted Regnase-1 KO cells and CD8 exhausted reference cells come from the same distribution. **D-E)** Top and side view of (predicted) CD8 exhausted cells from Regnase-1 KO in UMAP space, with ICA 25 embeddings on the z-axis.

**Supplementary Figure S9:**
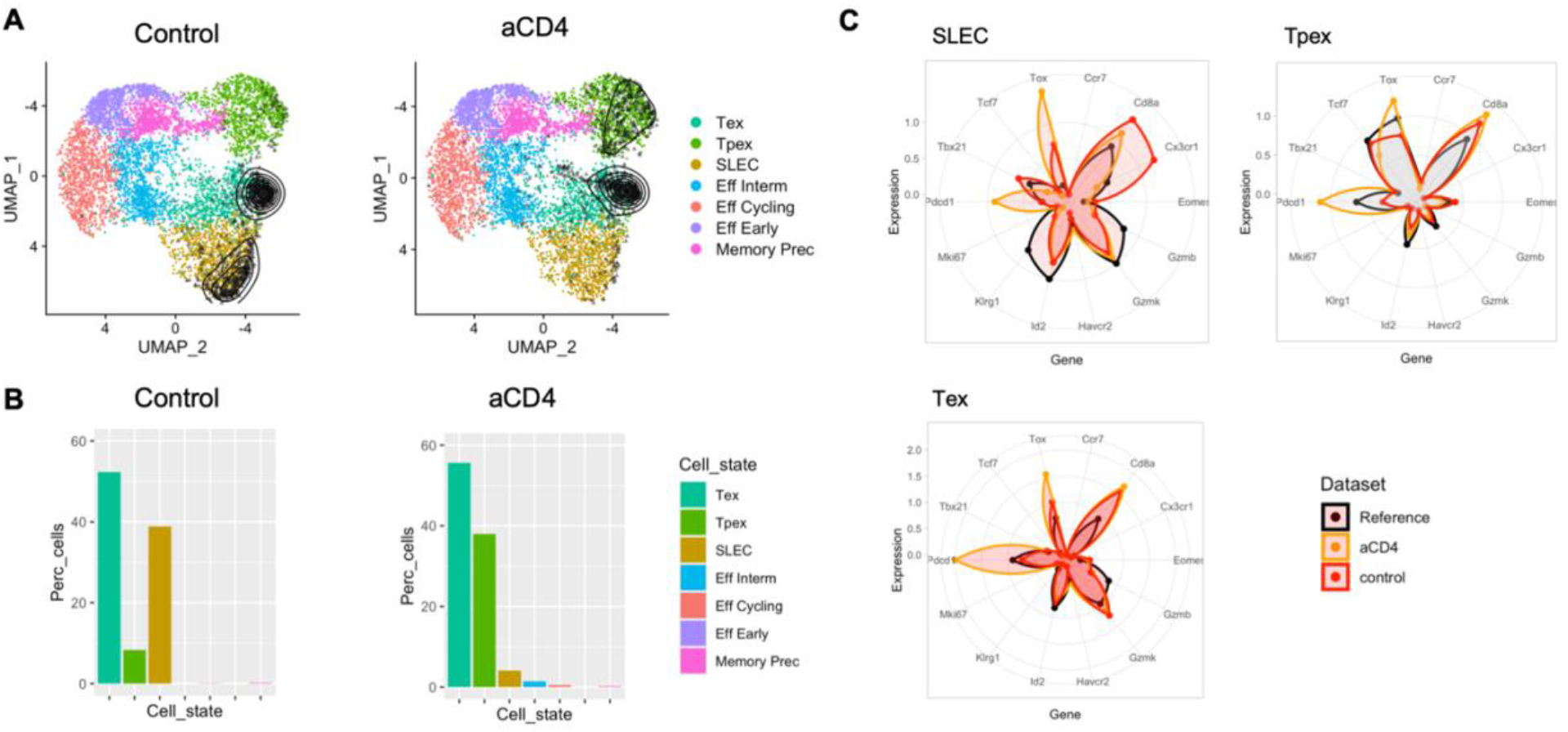
**A)** Projection of Control and CD4-depleted (aCD4) samples from Kanev *et al.* onto the LCMV reference atlas. **B)** Predicted cell subtype composition in terms of percentage of cells (Perc_cells) for Control and aCD4 treatment, showing a shift from short-lived effectors (SLEC) to Precursor Exhausted (Tpex) phenotype upon CD4 depletion. **C)** Normalized gene expression profile for selected marker genes for the SLEC, Tpex and Tex cell types.

**Supplementary Figure S10:**
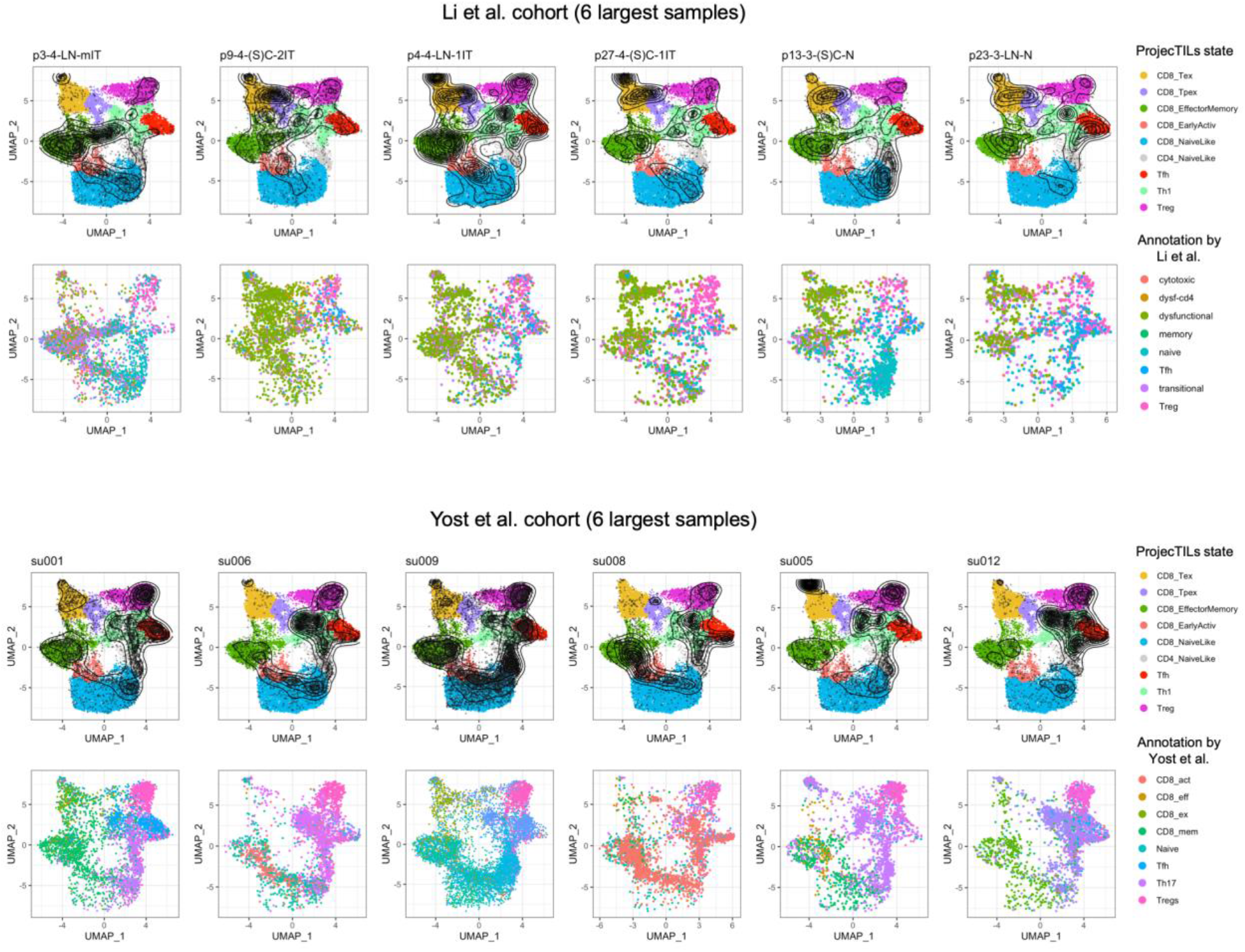
Projection of human scRNA-seq data onto the reference TIL atlas for 12 individual samples in two cancer patient cohorts. For the two cohorts (Li et al. and Yost et al.), each column represents a patient of the six with the largest number of cells. The top rows display the density of projected cells (as a contour line) over the reference murine TIL atlas, the bottom rows show individual projected cells colored by the original annotation by the authors. We note that patient su008 in the Yost et al. cohort has a disproportionate amount of cells annotated by the authors as Activated T cells (CD8_act) and projected in different sectors of the reference map; indeed, this subtype appears to be defined nearly completely by patient su008 (see Yost et al. (2019) Nature Medicine, Figure 2a), suggesting that the original definition of the Activated T cell subtype is largely explained by uncorrected batch effects.

**Supplementary Figure S11:**
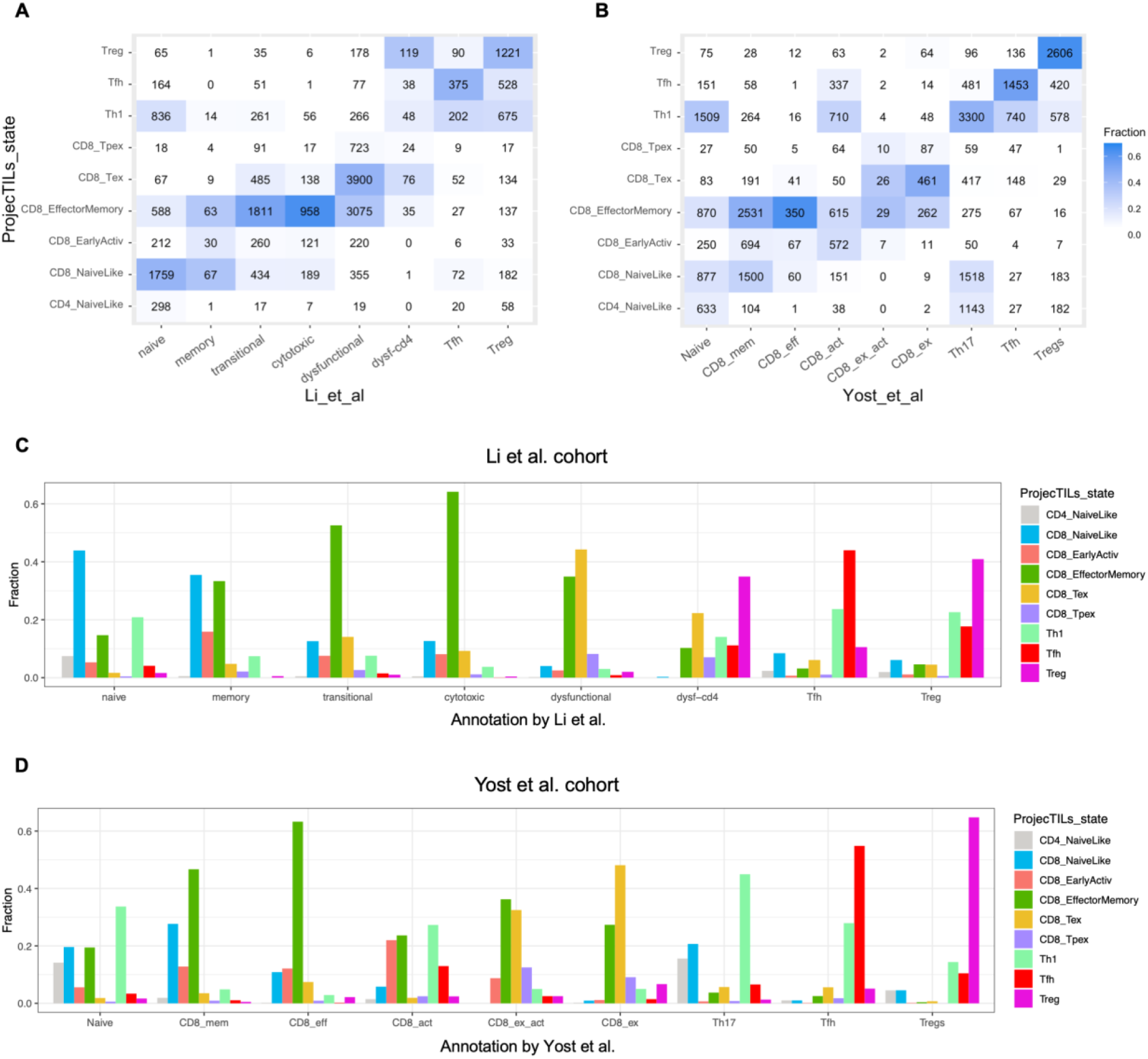
Concordance of ProjecTILs classification with cell annotation from original studies. **A)** ProjecTILs predicted states vs. original annotation for human T cell scRNA-seq data from Li *et al.*; numbers indicate absolute amount of cells for each annotation pair, colours indicate fraction over the total number in the original study annotation. **B)** Same as previous panel, but using the data and annotations from Yost *et al.* **C)** For each T cell type defined by Li *et al.*, fraction of cell assigned by ProjecTILs to the cell states of the mouse reference TIL atlas. **D)** For each T cell type defined by Yost *et al.*, fraction of cell assigned by ProjecTILs to the cell states of the mouse reference TIL atlas.

**Supplementary Figure S12:**
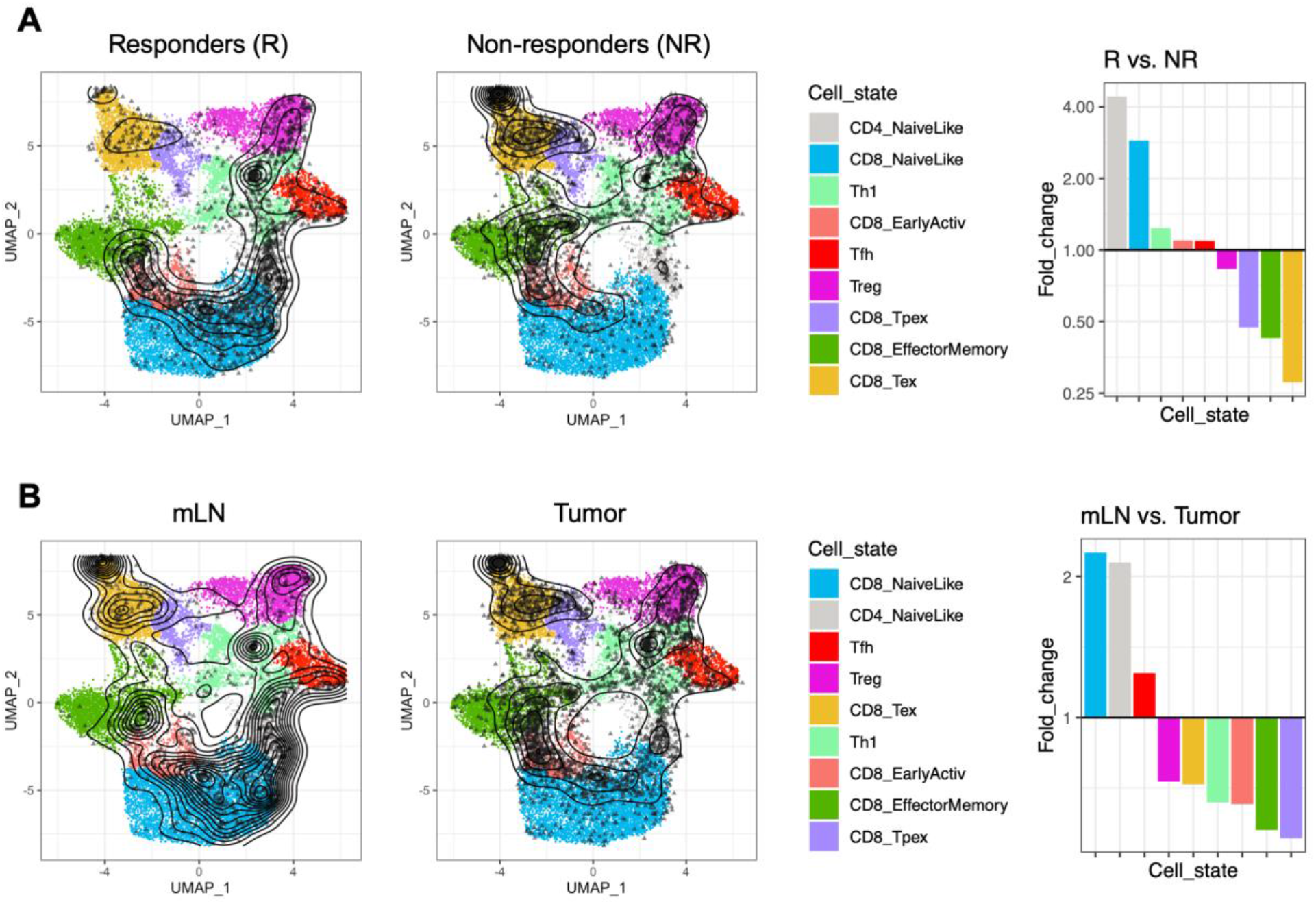
ProjecTILs analysis of baseline scRNA-seq data from melanoma patients (Sade-Feldman et al.). **A)** Distribution of projected query cells by response to immune checkpoint blockade, and relative enrichment of reference murine TIL states in responders (R) vs. non-responders (NR). **B)** Distribution of projected query cells by biopsy site (mLN: metastatic lymphnode vs. tumor), and relative enrichment of reference murine TIL states in the two tissues.

